# Epidermal Growth Factor Receptor Inhibition Prevents Caveolin-1-dependent Calcifying Extracellular Vesicle Biogenesis

**DOI:** 10.1101/2021.11.08.467799

**Authors:** Amirala Bakhshian Nik, Hooi Hooi Ng, Patrick Sun, Francesco Iacoviello, Paul R. Shearing, Sergio Bertazzo, Deniel Mero, Bohdan B. Khomtchouk, Joshua D. Hutcheson

**Affiliations:** Department of Biomedical Engineering, Florida International University, Miami, FL 33174, USA; Department of Human and Molecular Genetics, Herbert Wertheim College of Medicine, Florida International University, Miami, FL 33199, USA; The College at the University of Chicago, Chicago, IL 60637, USA; Department of Chemical Engineering, University College London, London WC1E 7JE, UK; Department of Medical Physics and Biomedical Engineering, University College London, London WC1E 6BT, UK; Dock Therapeutics Inc., Middletown, DE 19709, USA; Department of Medicine, Section of Computational Biomedicine and Biomedical Data Science, Institute for Genomics and Systems Biology, University of Chicago, Chicago, IL 60637, USA; Biomolecular Sciences Institute, Florida International University, Miami, FL 33199, USA

**Keywords:** Chronic Kidney Disease, Caveolin-1, Epidermal Growth Factor Receptor, Vascular Calcification, Extracellular Vesicles, Cardioinformatics, Drug Discovery

## Abstract

Chronic kidney disease (CKD) increases the risk of cardiovascular disease, including vascular calcification, leading to higher mortality. Release of calcifying extracellular vesicles (EVs) by vascular smooth muscle cells (VSMCs) promotes the ectopic mineralization of vessel walls. Caveolin-1 (CAV1) protein plays a key role in genesis of calcifying EVs in VSMCs. Epidermal growth factor receptor (EGFR) co-localizes with and influences the intracellular trafficking of CAV1. Using a diet-induced mouse model of CKD, we measured serum EGFR and assessed the potential of EGFR inhibition to prevent vascular calcification. Mice with CKD developed widespread vascular calcification, which associated with increased serum levels of EGFR. We computationally analyzed 7651 individuals in the Multi-Ethnic Study of Atherosclerosis (MESA) and Framingham cohorts to assess potential correlations between coronary artery calcium and single nucleotide polymorphisms (SNPs) associated with elevated serum levels of EGFR. Individuals in the MESA and Framingham cohorts with SNPs associated with increased serum EGFR exhibit elevated coronary artery calcium. In both the CKD mice and human VSMC culture, EGFR inhibition significantly reduced vascular calcification by mitigating the release of CAV1-positive calcifying EVs. EGFR inhibition also increased bone mineral density in CKD mice. Given that EGFR inhibitors exhibit clinical safety and efficacy in other pathologies, the current data suggest that EGFR may be an ideal target to prevent pathological vascular calcification.

## Introduction

Medial calcinosis manifests as the formation of calcium phosphate mineral in the media layer of arterial walls, leading to vascular stiffening, dysfunction, and cardiac overload [1, 2]. Medial calcinosis highly correlates with cardiovascular morbidity and mortality [3]. Calcification of arterial media commonly occurs in patients with chronic kidney disease (CKD) [2, 4]. CKD patients with no detectable vascular calcification have 8-year all-cause survival rates of around 90% compared to 50% survivability in age-matched patients with medial calcification [5]. Imbalanced serum calcium and phosphorous levels elevate the risk of medial calcinosis in CKD patients. Impaired renal excretion of phosphorous also leads to abnormal bone remodeling and mediates osteogenic differentiation of vascular smooth muscle cells (VSMCs) in the arterial walls [6].

Osteogenic differentiation of resident VSMCs and release of calcifying extracellular vesicles (EVs) mediate nucleation and growth of ectopic vascular calcification [3, 7]. This process mimics aspects of the physiological mineralization of osteoblasts and chondrocytes in bone via release of matrix vesicles [8]. Although calcifying EVs released into the vascular wall and bone matrix vesicles contribute to similar endpoints of mineralization, they originate through different pathways [9, 10]. The development of pharmaceuticals for vascular calcification targeting mechanisms specific to vascular calcifying EVs could avoid deleterious off-target effects on bone. Formation of calcifying EVs by VSMCs requires caveolin-1 (CAV1), a scaffolding membrane protein [11]. CAV1 resides in caveolar domains, small invaginations (50-100 nm) on the plasma membrane, which consist of the caveolin protein family, cholesterol, sphingolipids, and receptors [12, 13]. Caveolar functions include intra/extracellular lipid transfer, endocytosis, mechanotransduction, and signaling mediation [13, 14]. Calcifying VSMCs release CAV1-enriched EVs, and CAV1 knockdown abrogates calcification in these cells [11].

Epidermal growth factor receptor (EGFR) is a tyrosine kinase transmembrane glycoprotein [15], which localizes abundantly in caveolar domains. EGFR interacts with and modulates CAV1 trafficking [16] and recruits signaling proteins to caveolar domains [17]. EGFR actively participates in human cancer progression, and EGFR tyrosine kinase inhibition has become a widely utilized strategy in cancer therapies [18]. Both CAV1 and EGFR are elevated during breast cancer progression [19], and clinical studies indicate that overexpression of EGFR family members in breast cancer associates with increased ectopic calcification [20]. In cardiovascular pathogenesis, elevated EGFR activity correlates with oxidative stress and chronic inflammation [21]. EGFR inhibition in apolipoprotein E-deficient mice fed a high-fat diet prevented atherosclerotic plaque development [21]. However, the role of EGFR in VSMC-mediated calcification has not been reported.

Given these associations and the known interactions between CAV1 and EGFR, we hypothesized that EGFR inhibition would prevent vascular calcification by mitigating the biogenesis of calcifying EVs. We showed that EGFR inhibition reduces the release of pro-calcific CAV1-positive EVs and prevents calcification in osteogenic VSMC cultures and in CKD mice fed a high-phosphate diet. Furthermore, we computationally analyzed 7651 individuals in the Multi-Ethnic Study of Atherosclerosis (MESA) and Framingham cohorts, revealing a positive correlation between genetically mediated serum EGFR and coronary artery calcification (CAC) measured by computed tomography. Interestingly, EGFR inhibitor treatment also significantly reversed bone mineral loss in the CKD mice. Given the demonstrated clinical safety, our data suggest that EGFR inhibition could represent a viable therapeutic strategy to prevent vascular calcification in patients with CKD.

## Methods

### Chronic Kidney Disease and Vascular Calcification Mouse Model

The *in vivo* study was approved by the Institutional Animal Care and Use Committee (IACUC) at Florida International University under protocol AN20-006 and conformed to current NIH guidelines. The experimental design was based upon procedures established in a previous study to induce CKD and vascular calcification in mice [22]. 8-week-old wild type C57BL/6J mice (n = 38, 19 per biological sex) were fed an adenine-supplemented diet (0.2%, TestDiet, Richmond, IN) for 6 weeks to induce severe kidney injury. The mice then received a diet containing 1.8% phosphate (TestDiet, Richmond, IN) and 0.2% adenine for an additional two weeks to induce medial calcinosis. Along with this calcifying diet, a group of mice (n = 19) received daily tyrphostin AG1478 (10 mg/kg mouse, Millipore Sigma, T4182) via oral gavage. The remaining mice (n = 19) received vehicle treatment (1% w/v, carboxymethylcellulose sodium salt, Sigma, C5678). For non-diseased controls, a third group of mice (n = 12, 6 per biological sex) were fed a regular chow diet and received the vehicle for the final two weeks. During the oral gavage, animals were partially anesthetized using isoflurane (1%, Patterson Veterinary, 07-893-1389, in 2 L.min^-1^ oxygen flow). All animals received a tail vein injection with the calcium tracer OsteoSense 680EX (80 nmol/kg mouse, PerkinElmer, NEV10020EX) 48 hours prior to euthanization. At study endpoint, mice were anesthetized with isoflurane (1%, in 2 L.min^-1^ oxygen flow) followed by retro-orbital bleeding for blood collection. Mice were then immediately euthanized by laceration of the diaphragm before tissue collection. After resection, the aortas were imaged using a near-infrared scanner (LI-COR Odyssey) to visualize the vascular calcification burden. A custom MATLAB script quantified the total area of the calcium tracer, which was normalized to the total scanned aorta area.

Immediately after scanning, the tissue was incubated in a digestive solution [23] of sucrose (0.25 M, Sigma, S7903), NaCl (0.12 M, Fisher Chemical, BP358), KCl (0.01 M, Fisher Chemical, P217), Tris hydrochloride (0.02 M, Fisher Chemical, BP153), and collagenase (600 U/mL, Worthington Biochemical, LS004174) for 2 hours at 37°C. The solution was then centrifuged at 1,000×g for 15 min to remove cell debris and at 33,000×g for 30 min to remove microvesicles. Finally, the supernatants were ultracentrifuged (Beckman Coulter, Optima MAX-TL) at 100,000×g for 1 hour to isolate the EVs of interest. The pellet was suspended in RIPA lysis and extraction buffer (G Biosciences, 786-489) supplemented with pierce protease inhibitor (Thermo Scientific, A32963). To yield sufficient protein concentration for analysis, EVs isolated from 2 to 3 aortas were pooled.

### Osteogenic Stimulation, In vitro Calcification, and Extracellular Vesicle Isolation

Primary human coronary artery vascular smooth muscle cells (VSMCs, ATCC, PCS-100-021) were cultured using vascular smooth muscle cell media and growth kit (ATCC, PCS-100-042). VSMCs (passage 4-6) were harvested using 0.05% trypsin-EDTA solution (Caisson Labs, TRL04) and seeded with a density of 26,320 cells.cm^-2^ and incubated for 72 hours at 37°C, 5% CO_2_ with controlled humidity prior to treatment. VSMCs were treated with either control media, consisting of DMEM (HyClone, SH30022.01), 10% v/v bovine calf serum (iron-supplemented, R&D Systems, S11950), and 1% v/v penicillin-streptomycin (Gibco, 15070-063), or with an osteogenic media (OS) optimized to induce calcification [24, 25]. OS media were supplemented with 10 mM β-glycerophosphate (Sigma, 13408-09-8), 0.1 mM L-ascorbic acid (Sigma, 113170-55-1), and 10 nM dexamethasone (Sigma, 50-02-2). To assess the role of EGFR inhibition, tyrphostin AG1478 (Millipore Sigma, T4182) was dissolved in the vehicle (DMSO:Methanol, 1:1) and added to OS media to a final concentration of 2.5 μM. An equal volume of the vehicle was added to the control and OS groups. We found that 28 days in OS culture media led to robust calcification by VSMCs; therefore, all cultures (n = 3, independent donors, male and female) were treated for 28 days and media were replaced every three days. On days 6, 13, 20, and 27 the media were replaced by an extracellular-vesicle-free (EV-free) media (ultracentrifuged for 15 hours at 100,000×g at 4°C to remove background EVs common in the serum). After 24 hours, conditioned media were collected on days 7, 14, 21, and 28. Collected media were centrifuged at 1,000×g for 5 min to remove cell debris. EV isolation was performed using ultracentrifugation at 100,000×g for 1 hour.

Osteoblasts (from human fetus, hFOB 1.19, ATCC, CRL-11372) were cultured and grown in DMEM containing 10% v/v bovine calf serum and 1% v/v penicillin-streptomycin. Osteoblasts (passage 4-6) were harvested using 0.25% trypsin-EDTA solution (Caisson Labs, TRL01), seeded with a density of 5,200 cell.cm^-2^, and incubated for 24 hours at 37°C and 5% CO_2_ with controlled humidity. The cells were treated in three groups of control, OS, and OS supplemented with tyrphostin AG1478 (2.5 μM) for 21 days and media were changed every three days. Compared to VSMCs, we observed more rapid mineralization in osteoblasts cultured in OS with full matrix mineralization apparent after 21 days. Similar to the VSMC experiments, EV-free media were added to the cultures on days 6, 13, and 20, and collected 24 hours later on days 7, 14, and 21. Collected media were centrifuged at 1,000×g for 5 min to remove cell debris. Matrix vesicles were isolated using the ultracentrifugation at 100,000×g for 1 hour.

### Alizarin Red S Staining and Quantification

At the end of experiments (28 and 21 days of treatment for VSMCs and osteoblasts, respectively), media were removed, and the cells were fixed using formalin (10%, Fisher Chemical, SF100) for 15 min. To visualize *in vitro* calcification, Alizarin Red S stain (ARS, Ricca, 500-32) was added to the wells and incubated for 30 min at room temperature. The stain was then removed, and the cells were washed three times with milliQ water. To quantify the *in vitro* calcification, ARS stain was extracted using acetic acid (1.67 M, Fisher Chemical, A38S) on a shaker. After 30 min, the supernatants were collected, briefly vortexed, and heated at 85°C for 10 min. The samples were then cooled on ice for 5 min and centrifuged at 20,000×g for 15 min to remove background particles. Sample absorbance of 405 nm light was measured using a multi-mode reader (BioTek, Synergy HTX).

### Kidney Histological Analysis

To assess histological changes in kidneys due to renal injury, Hematoxylin and Eosin (H&E) staining was performed. The kidneys resected from the mice were fixed using formalin (10%) for three hours. Tissues were embedded using Tissue-Plus OCT (Fisher Scientific, 23-730-571). The samples were cryosectioned with a thickness of 12 μm and stained using rapid chrome H&E staining kit (Thermo Scientific, 9990001).

### Quantitative Real Time Polymerase Chain Reaction

Following 7 or 14 days in control, OS, or OS plus EGFR inhibitor media, VSMCs and osteoblasts were lysed in 1 mL TRIzol solution (Invitrogen, 15596018). Total RNA was isolated according to the manufacturer’s protocol. To perform the quantitative real time polymerase chain reaction (qRT-PCR), Power SYBR Green RNA-to-CT 1-Step Kit (Applied Biosystems, 4391178) was used. 50 ng of isolated template RNA were added to each reaction for qRT-PCR. The results were normalized to Glyceraldehyde 3-phosphate dehydrogenase (GAPDH) expression level as the housekeeping control. The relative gene expression levels were calculated using comparative CT method, considering control groups as the reference. The following human primers were purchased from Integrated DNA Technologies (IDT); *GAPDH* Forward: CTTCGCTCTCTGCTCCTCCTGTTCG and Reverse: ACCAGGCGCCCAATACGACCAAAT; *RUNX2* Forward: GCTCTCTAACCACAGTCTATGC and Reverse: AGGCTGTTTGATGCCATAGT; *ALPL* Forward: GGAGTATGAGAGTGACGAGAAAG and Reverse: GAAGTGGGAGTGCTTGTATCT; *Osteocalcin (BGLAP)* Forward: TCACACTCCTCGCCCTATT and Reverse: CCTCCTGCTTGGACACAAAA.

To isolate RNA from the resected kidneys, the tissues were homogenized using a grinder (Sigma, Z529672) and lysed in 1 mL TRIzol solution. After 10 min incubation at room temperature, the samples were centrifuged at 12,000×g for 10 min at 4°C. The supernatants were collected and 200 μL of chloroform (Sigma Aldrich, C2432) were added to each sample. The samples were vortexed, incubated at room temperature for 10 min, and centrifuged for 15 min at 12,000×g, 4°C. The aqueous phase was collected from each sample and 500 μL of isopropanol were added; the samples were vortexed, incubated for 15 min at room temperature followed by 15 min on ice, and centrifuged at 21,000×g for 15 min. The supernatants were discarded, and pellets were washed twice with 500 μL cold ethanol (75% v/v) and centrifuged at 21,000×g for 5 min [26–28]. The isolated RNA templates were heated at 65°C for 15 min, and the concentrations were measured using a spectrophotometer (NanoDrop Lite, Thermo Scientific). Power SYBR Green RNA-to-CT 1-Step Kit with 100 ng isolated template RNA per reaction was used. The following mouse primers were purchased from Eurofins Scientific; *Gapdh* Forward: AACGACCCCTTCATTGAC and Reverse: TCCACGACATACTCAGCAC; *Col1a1* Forward: CCTCAGGGTATTGCTGGACAAC and Reverse: ACCACTTGATCCAGAAGGACCTT; *Tgfb1* Forward: TGGAGCAACATGTGGAACTC and Reverse: CAGCAGCCGGTTACCAAG.

### Alkaline Phosphatase Activity Assay

To assess the activity of intracellular tissue non-specific alkaline phosphatase (TNAP), a colorimetric assay kit (BioVision, K412) was used. VSMCs (n = 3) after 14 days and osteoblasts (n = 3) after 7 days, were lysed in 120 μL assay buffer. 80 μL of each sample were mixed with 50 μL of 5 mM pNPP solution and incubated for 60 min at 25°C. The colorimetric change resulting from the reaction was detected using a plate reader to measure absorbance at 405 nm. The results were normalized to the total protein for associated samples measured by a BCA protein assay (BioVision, K813). For EV or matrix vesicle TNAP activity measurement, after ultracentrifugation at 100,000×g for 1 hour, the pellets were re-suspended in 120 μL assay buffer. The assessment was performed using the same assay protocol described for intracellular TNAP activity and the results were normalized to the total protein for each sample. For mouse serum TNAP activity, the samples were diluted 1:20 and assessed according to the manufacturer’s protocol.

### Serum Creatinine and Urea Nitrogen Assessment

To measure serum creatinine, a colorimetric assay (Cayman Chemicals, 700460) was used; 15 μL of each collected serum were added to 200 μL of a solution of assay reaction buffer and color reagent (1:1), incubated for 1 min at room temperature and measure at 495 nm using a multi-mode reader. The absorbance was measured at 495 nm for second time after 7 min. The changes in optical density (ΔO.D.) for each sample were associated to the creatinine concentration according to the manufacturer’s protocol.

To assess serum urea nitrogen, the serum samples were diluted 1:10; a colorimetric assay (Invitrogen, EIABUN) measured the serum urea nitrogen. Briefly, collected serum (50 μL) was mixed with 150 μL assay color solution (reagent A: reagent B, 1:1) and incubated at room temperature for 30 min. the colorimetric changes were measure at 450 nm.

### Extracellular Collagen Assessment

After 28 days of treatment, soluble collagen was extracted from the cultures using acetic acid (0.5 M) through overnight incubation at 4°C. A colorimetric assay, Sircol soluble collagen assay (Biocolor, S1000), measured the total soluble extracellular matrix (ECM) collagen in each group. Samples were prepared and assessed according to the manufacturer’s protocol. Results were then normalized to the total protein measured using BCA assay.

### Subcellular Protein Fractionation for VSMCs and Aortas

8-week-old wild type C57BL/6J mice (n = 20, female) received the adenine-supplemented diet for 6 weeks to induce CKD, followed by two additional weeks of the diet containing 1.8% phosphate and 0.2% adenine to induce medial calcinosis. Mice were split into two groups (10 per group). The first group received daily tyrphostin AG1478 (10 mg/kg mouse), while the other group received vehicle (1% w/v, carboxymethylcellulose sodium salt). At study endpoint, the animals were euthanized, and the aortas were resected. A subcellular protein fractionation kit for tissue (Thermofisher, 87790) was used to isolate cellular cytosolic fraction from the resected aortas, using the manufacturer’s protocol. Briefly, the tissues were minced and homogenized using a grinder. The samples were then incubated in a cytoplasmic extraction buffer for 10 min at 4°C, followed by centrifugation at 1000×g for 5 min. The supernatants yielded the cytosolic fraction. To obtain sufficient protein for analyses, two aortas were pooled per data point.

VSMCs were treated with control, OS, and OS supplemented with tyrphostin AG1478 (2.5 μM) for 14 days. At the experiment endpoint, using a subcellular protein fraction kit for cultured cells (Thermofisher, 78840), cytosolic fraction was isolated according to the manufacturer’s protocol. Briefly, VSMCs were harvested using 0.25% trypsin solution and resuspended in cytoplasmic extraction buffer. After 10 min incubation at 4°C, the samples were centrifuged at 1000×g for 5 min and the supernatants were collected as cytosolic fractions. The protein concentration for aortic tissue and VSMC fractions were quantified using a BCA assay and samples were prepared for protein immunoblotting.

### Immunoprecipitation and Lipid Raft Isolation

Following 14 days of treatment, VSMCs (n = 3) were lysed using Pierce immunoprecipitation lysis buffer (Thermo Scientific, 87788) supplemented with protease inhibitor. A Dynabeads Protein G immunoprecipitation kit (Invitrogen, 10007D) was used to precipitate CAV1 from the cell lysates [11]. Briefly, Dynabeads coated with protein G were incubated with either CAV1 antibody (5 μg, abcam, ab17052) or IgG mouse control antibody (5 μg, Proteintech, B900620) by rotation for 3 hours at 4°C. After removal of supernatants using a magnet, 100 μg of protein were loaded to the beads and incubated for 5 hours at 4°C while rotating. After removal of supernatants and 3 washes with washing buffer, 40 μL of elution buffer (from the kit) and 20 μL of 1:1 NuPAGE sample reducing agent (Thermo Scientific, NP0009) and NuPAGE LDS sample buffer (Thermo Fisher Scientific, NP0007) were added to the beads. The samples were incubated by rotation at 4°C for 30 min, and denatured at 70°C for 10 min. After removal of beads by a magnet, the samples were ready for protein immunoblotting.

To isolate lipid rafts, VSMCs (n = 3) were treated under control, control plus EGFR inhibitor, OS, or OS plus EGFR inhibitor for 14 days. Then, the cells were lysed in a buffer containing HEPES (25 mM, Fisher Scientific, BP310-100), NaCl (150 mM), PMSF (1 mM, Boston Bioproducts, PI120), EDTA (1 mM, Invitrogen, AM9260G), Triton X-100 (1% v/v, Fisher Scientific, BP151-100), and protease inhibitor. Five gradient layers were prepared using OptiPrep density gradient medium (Sigma-Aldrich, D1556), including 35% (with cell lysates), 30%, 25%, 20%, and 0% (with lysis buffer), and loaded to ultracentrifuge tubes (Optiseal bell, Beckman Coulter, 361621), respectively (total volume of 4.5 mL). Samples were ultracentrifuged at 4°C and 110,000×g for 4 hours; 9 fractions (500 μL per fraction) were isolated for each group and used for protein immunoblotting.

### Gel Electrophoresis and Protein Immunoblotting

VSMCs, osteoblasts, isolated EVs (either from cells or mouse aortas), and matrix vesicles (from osteoblasts) were lysed in RIPA lysis and extraction buffer supplemented with protease inhibitor. After adding Laemmli SDS-sample buffer (1:4 v/v, Boston BioProducts, BP-110R) to each lysate, the samples were denatured at 100°C for 10 min, loaded into 7.5-12% 1-mm SDS-PAGE gel (15 to 20 μg protein per lane), and run at 170 V. The proteins were then transferred to Trans-Blot turbo nitrocellulose membranes (BIO-RAD, 1704158) at 25 V for 7 min. To quantify the total protein, the membranes were stained using 2% w/v Ponceau stain (Alfa Aesar, AAJ6074409) for 20 min, followed by one wash with 5% acetic acid and milliQ water for 5 min. After imaging, the intensity of each lane was measured in ImageJ for total protein normalization. Membranes were blocked with 5% w/v bovine serum albumin (HyClone, SH30574.01) in TBS-Tween (1X) for 1 hour. The membranes were incubated with primary antibodies of interest, including CAV1 (1:200, Abcam, ab2910), TNAP (1:200, Invitrogen, 702454), EGFR (1:00, EMD Millipore, 06-874), CD63 (1:200, Abcam, ab231975), GAPDH (1:100, Abcam, ab181602), and Annexin V (1:200, proteintech, 11060-1-AP) overnight at 4°C. After three washes with TBS-Tween (1X), the membranes were incubated with secondary antibody (1:1000, Li-Cor) for 1 hour, followed by three washes with TBS-Tween (1X). The protein bands were visualized with Odyssey CLx scanner (Li-Cor) and quantified using Image Studio Lite software (Li-Cor). All western blotting images and corresponding Ponceau stains used for normalization are provided in **Online materials, Fig. I and II**.

### Immunofluorescence Staining and Imaging

VSMCs were fixed after 14 days of culture using formalin (10%) for 15 min and washed with PBS. A solution of PBS and Triton X (0.1% v/v) permeabilized the plasma membrane for 10 min at room temperature. To avoid non-specific antibody binding, the cells were incubated with a blocking buffer solution, consisting of BSA (1% w/v) and glycine (22.5 mg/mL) in PBS for 30 min at room temperature. Next, the cells were incubated for 2 hours with primary antibody against CAV1 (1:200) and washed three times with PBS. Cells were then incubated with a secondary antibody, Alexa Fluor 594 (1:500, Abcam, ab150080), for 1 hour at room temperature, followed by three washes with PBS. To visualize actin filaments, samples were incubated for 20 min with Phalloidin-iFluor 488 conjugate (1:50, Cayman Chemical, 20549) followed by three washes with PBS.

Resected mouse aortas were fixed in formalin (10%) for 2 hours. The tissues were rinsed with PBS and embedded in OCT. The samples were cryosectioned with a thickness of 7 μm. The samples were incubated with a blocking buffer containing donkey serum (10% v/v), Triton X (0.3% v/v), and BSA (1% w/v) in PBS for 1 hour at room temperature. After blocking buffer removal, a solution of donkey serum (1% v/v), Triton X (0.3% v/v), BSA (1% w/v) in PBS, with primary antibody against either CAV1(1:200), EGFR (1:100), or TNAP (1:200) was added to the samples. After an hour incubation at room temperature, the primary antibody solution was removed, and the samples were washed with PBS. Secondary antibody, Alexa Fluor 594 (1:500, Invitrogen, A21207) was added to the samples and incubated for 1 hour at room temperature. After washing the samples with PBS, they were stained with DAPI (0.2 μg/mL, Cayman Chemical, 14285) for 10 min and washed with PBS. The samples were mounted using Flouromount (Sigma Millipore, F4680). A confocal microscopy system (Eclipse Ti, Nikon) was used to image both cellular and tissue samples.

### X-ray Computed Tomography (X-ray CT)

Femurs were dissected from mice, wrapped in parafilm and imaged directly in a Nikon XT H 225 scanner (macro-CT, Nikon Metrology, Tring, UK). The raw transmission images were reconstructed using commercial image reconstruction software package (CT Pro 3D, Nikon Metrology, Tring, UK), which employs a filtered back-projection algorithm. The scan was performed using 80 kV beam energy, 70 μA beam current, and a power of 5.6 W. A PerkinElmer 1620 flat panel detector was used, with 200 μm pixel size. The resulting effective pixel size was 5 μm. The exposure time per projection was 0.5 s, and a total of 1601 projections were acquired, resulting in a scanning time of approximately 13 minutes per sample. Bone structural parameters, including thickness and volume fraction (the ratio of bone volume (BV) to total volume (TV)), for both cortical and trabecular regions were assessed using a plug-in module, BoneJ, in ImageJ (NIH, USA) [29].

### Identification of Instrumental Variables for Mendelian Randomization

Instrumental variables (IVs) were selected using an agnostic p-value threshold, p < 5×10^-6^, as advised by the methodological literature on Mendelian Randomization (MR) [30]. Single nucleotide polymorphisms (SNPs) associated with significantly elevated serum EGFR concentration (p < 5×10^-6^) from a previous proteomics study were compared against genotyped SNPs in the Multi-Ethnic Study of Atherosclerosis (MESA) SNP Health Association Resource (SHARe), and all SNPs presented in both the proteomics study and MESA genotyping data associated beyond this p-value threshold were included as IVs for the MR analysis [31]. In total, three SNPs of rs12666347, rs2371816, and rs7806938 were included. The same 3 IVs and measure of CAC were used to replicate the significance of the MR analysis and validate results in the Offspring Cohort of the Framingham Heart Study (FHS).

### Calculation of SNP-EGFR and SNP-CAC Association in the MESA and FHS Cohorts

Effect sizes of each SNP on EGFR concentration, as well as their standard errors, were extracted from the publicly available summary statistics [31]. To calculate the effect sizes of each SNP on calcification levels, we identified 1,896 individuals from the FHS Offspring cohort, and 5,755 individuals who completed MESA Exam 1 who had available genotyping information. For each of these individuals, genotyping information, age, sex, study site, race, and Agatston score were extracted. Agatston scores are a measure of CAC determined through cardiac imaging, with an increasing Agatston score representing increased CAC. Associations between each IV SNP and CAC is calculated using logistic regression, treating Agatston scores as a binary variable (= 0 vs > 0) and including age, sex, study site, and race as covariates in the model. All analyses were conducted using the R programming language.

### Mendelian Randomization

Following identification of SNP-CAC and SNP-EGFR association and standard error values, MR analysis was performed to determine the presence and estimate the magnitude of causal effect that elevated serum EGFR has on CAC. 11 different regressions were included in the MR analysis to correct for possible pleiotropic effects, a possible source of confounding. Included regressions were simple median, weighted median, penalized median, inverse-variance weighted (IVW), penalized IVW, robust IVW, penalized-robust IVW, MR-Egger, penalized MR-Egger, robust MR-Egger, and penalized-robust MR-Egger. MR analysis was performed using the Mendelian Randomization package in R [32, 33] (R Core Team, 2021; Yavorska and Staley, 2021). We accounted for multiple testing errors using a Bonferroni-adjusted 0.05 significance level of 0.0045 (0.05/11).

### Statistics

Data are presented as the mean of independent replications, and error bars represent the standard error of the mean. The reported *n* values represent independent biological replicates. Statistical significance between groups was calculated using one-way ANOVA with Tukey’s post-hoc test in GraphPad Prism 8. A p-value less than 0.05 was considered statistically significant. In case of comparison between two groups, the statistical significance was calculated using t-test with p-values less than 0.05.

## Results

### EGFR inhibition reduces vascular calcification in a CKD mouse model

Visualization of the calcium tracer, OsteoSense, showed widespread vascular calcification in CKD mice compared to the chow-fed control group. Daily EGFR inhibitor gavage (10 mg/kg/mouse) for two weeks dramatically reduced vascular calcification in CKD animals (**Fig. 1, a**). Quantification of the OsteoSense intensity revealed a significant reduction in vascular calcification in the EGFR inhibited group (p ≤ 0.0001), as shown in **Fig. 1, b**. The level of serum EGFR was elevated in the CKD group compared to chow fed animals (p = 0.038), with no significant difference between CKD and EGFR inhibited groups (p = 0.78) (**Fig. 1, c**). Serum TNAP activity, urea nitrogen, and creatinine (**Fig. 1, d** to **f**) in CKD animals were significantly elevated compared to the control group (p = 0.003, p ≤ 0.0001, and p ≤ 0.0001, respectively). EGFR inhibition did not reduce serum TNAP activity (p = 0.06), urea nitrogen (p = 0.82), and creatinine (p = 0.94). Gene expression of common renal fibrosis markers, *Tgfb1* and *Col1a1* (**Fig. 1, g** and **h**), were significantly increased in both CKD mice (p = 0.02 and p = 0.02 for *Tgfb1* and *Col1a1*, respectively) and CKD mice treated with EGFR inhibitor (p = 0.02 and p = 0.03 for *Tgfb1* and *Col1a1*, respectively) when compared to chow-fed controls, with no significant differences between the CKD groups (p = 0.7 and p = 0.6 for *Tgfb1* and *Col1a1*, respectively). Qualitative assessment of histological sections of resected kidney tissues showed enlarged tubular structures in both CKD and EGFR inhibitor treated CKD groups, compared to the chow-fed control (**Fig. 1, j**). These results indicate that EGFR inhibition reduces vascular calcification in CKD animals independent of effects on renal injury.

**Figure 1.**
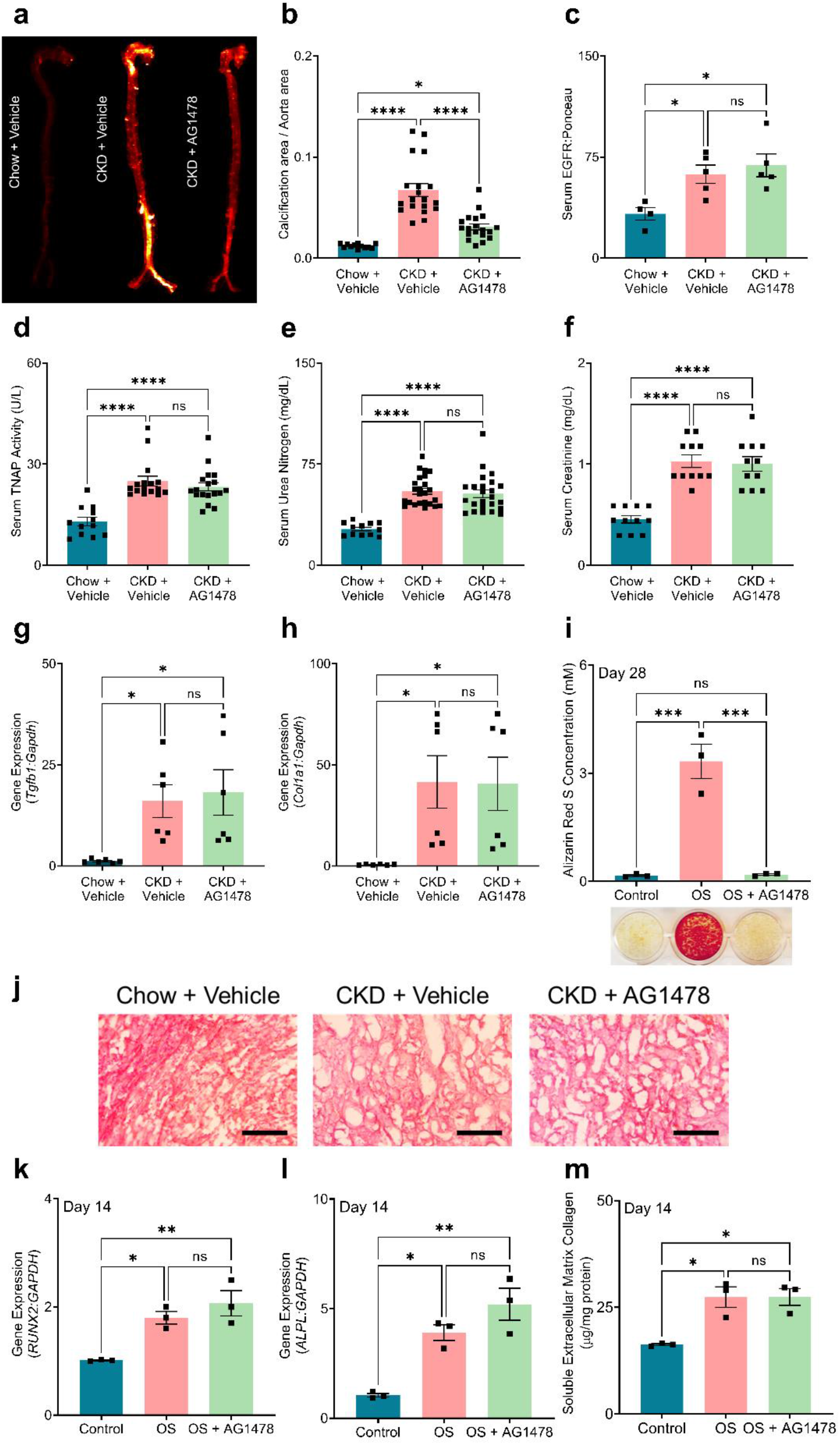
EGFR inhibition prevents vascular calcification *in vivo* and *in vitro*. (a) Visualization of vascular calcification using calcium tracer OsteoSense; (b) Quantification of the OsteoSense to correlate with vascular calcification burden (n = 50); (c) Serum EGFR level collected from mouse groups; (d) Serum TNAP activity collected from mouse groups; (e) Serum urea nitrogen level collected from mouse groups; (f) Serum creatinine level collected from mouse groups; (g and h) Gene expression of renal fibrotic markers, *Tgfb1* and *Col1a1*; (i) *In vitro* calcification visualization using Alizarin Red S staining and quantification; (j) H&E staining of mouse kidney tissues (20X, scale bar 0.5 mm); (k and l) Gene expression of osteogenic markers, *RUNX2* and *ALPL* in VSMCs following 14 days of treatment; (m) Extracellular matrix collagen accumulation in VSMC cultures. *P < 0.05, **P ≤ 0.01, ***P ≤ 0.001, and ****P ≤ 0.0001, ANOVA with Tukey’s post-hoc test.

### EGFR inhibition attenuates in vitro vascular smooth muscle cell calcification

VSMCs calcified following 28 days of culture in OS media, as shown by ARS staining (**Fig. 1, i**, representative image). Treatment of OS cultures with EGFR inhibitor abrogated *in vitro* calcification of the VSMCs **(Fig. 1, i**). Gene expression analysis of the common osteogenic markers, *RUNX2* and *ALPL*, revealed that VSMCs cultured in both OS (p = 0.02 and p = 0.02 for *RUNX2* and *ALPL*, respectively) and OS treated with EGFR inhibitor (p = 0.04 and p = 0.03 for *RUNX2* and *ALPL*, respectively) acquired an osteogenic phenotype after 14 days of culture (**Fig. 1, k** and **l**), with no significant differences between the groups (p = 0.4 and p = 0.1 for *RUNX2* and *ALPL*, respectively). Moreover, OS media promoted the accumulation of ECM collagen *in vitro*, which creates a platform for calcifying EVs to initiate calcification [25] (**Fig. 1, m**); EGFR inhibition did not affect the ECM collagen accumulation (p = 0.10). These data indicate that EGFR inhibition attenuates VSMC calcification without affecting VSMC phenotypic changes.

### EGFR inhibition alters CAV1/TNAP intracellular trafficking

Both OS cultured VSCMs and OS cultured VSMCs treated with EGFR inhibitor significantly increased the total level of intracellular CAV1 in VSMCs compared to the control group (p < 0.0001) (**Fig. 2, a**). OS media also increased intracellular EGFR in VSMCs compared to the control group (p = 0.019, **Fig. 2, b**). EGFR inhibition prevented the OS-induced increase in EGFR protein (p = 0.84). Similar to the gene expression data (**Fig. 1, l**), both OS cultured VSMCs and OS cultured VSMCs treated with EGFR inhibitor exhibited elevated intracellular TNAP activity (p = 0.03 and p = 0.03 for intracellular CAV1 and TNAP activity, respectively, compared to control) (**Fig. 2, c**). Confocal micrographs of VSMCs (**Fig. 2**, panel **d**) showed alignment of CAV1 protein along actin filaments in VSMCs cultured in OS media. In the OS cultured VSMCs treated with EGFR inhibitor, larger clusters of CAV1 were observed between filaments. Subcellular protein fractionation of VSMCs revealed that both intracellular CAV1 and TNAP were elevated in EGFR inhibited cultures compared to control (p = 0.02 and p = 0.003, respectively) and OS groups (p = 0.04 and p = 0.005, respectively, **Fig. 2, e** and **f**). Qualitative analysis of confocal micrographs of CAV1, EGFR, and TNAP immunofluorescence in the aorta of mice indicated elevation of all three proteins in CKD mice and CKD mice treated with EGFR inhibitor, compared to the chow-fed controls (**Fig. 3**, panel **a**). Subcellular protein fractionation of aorta indicated higher intracellular CAV1 and TNAP proteins in EGFR inhibited CKD animals compared to the CKD group (p = 0.04 and p = 0.0001, and p = 0.018, respectively), similar to *in vitro* data (**Fig. 3, b** to **d**).

**Figure 2.**
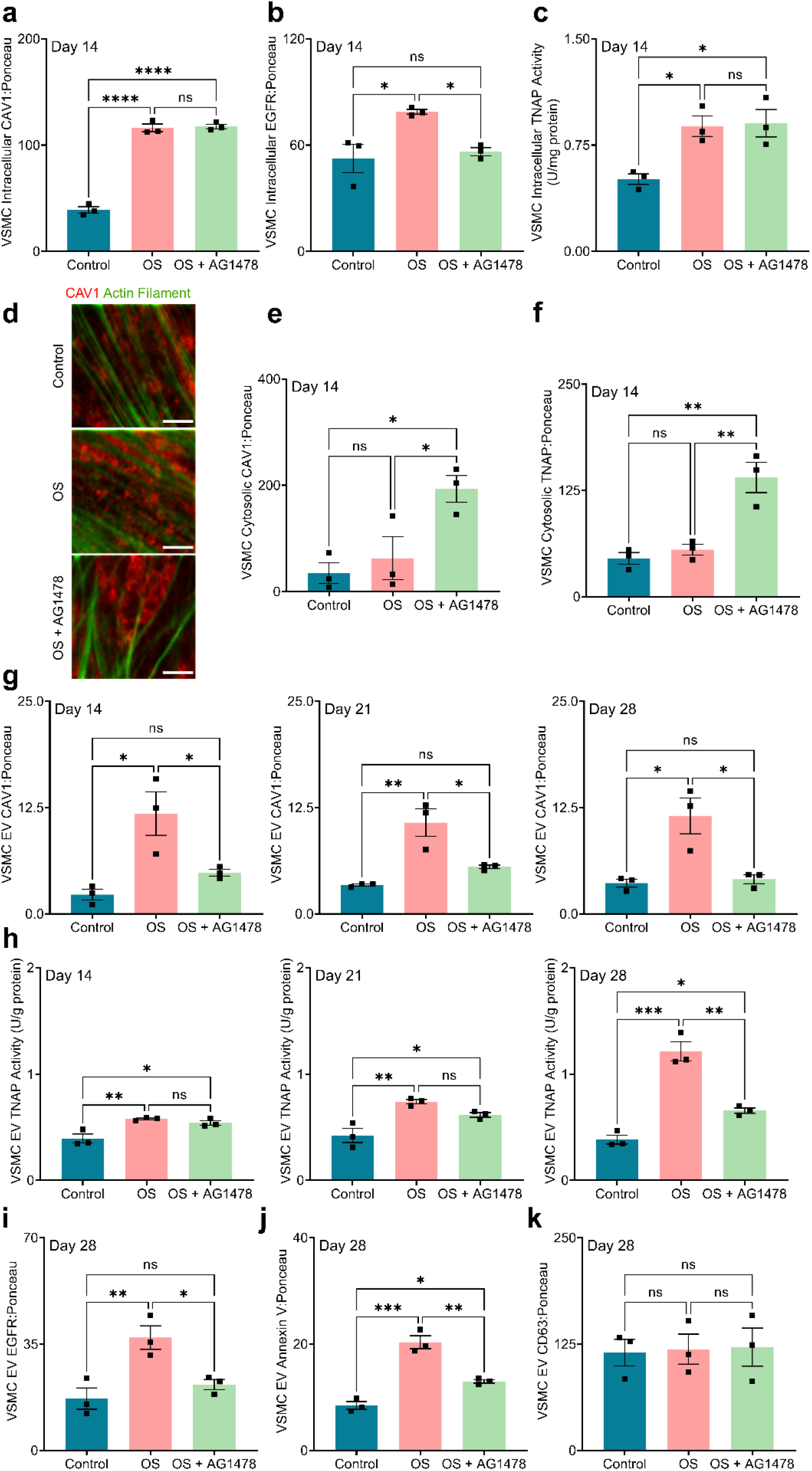
EGFR inhibition modulates CAV1 trafficking in VSMCs. Intracellular level of: (a) CAV1, (b) EGFR, and (c) TNAP activity in VSMCs after 14 days of culture; (d) Confocal micrographs of CAV1 distribution in VSMCs following 14 days of treatment (1200X, scale bar: 0.5 μm); Cytosolic level of: (e) CAV1, and (f) TNAP protein following 14 days of treatment; (g) CAV1 level on EVs isolated from VSMC cultures after 14, 21, and 28 days; (h) TNAP activity of the EVs isolated from VSMC cultures after 14, 21, and 28 days; EV level of: (i) EGFR, (j) Annexin V, and (k) CD63 liberated from VSMCs on day 28 of treatment. *P < 0.05, **P ≤ 0.01, ***P ≤ 0.001, and ****P ≤ 0.0001, ANOVA with Tukey’s post-hoc test.

**Figure 3.**
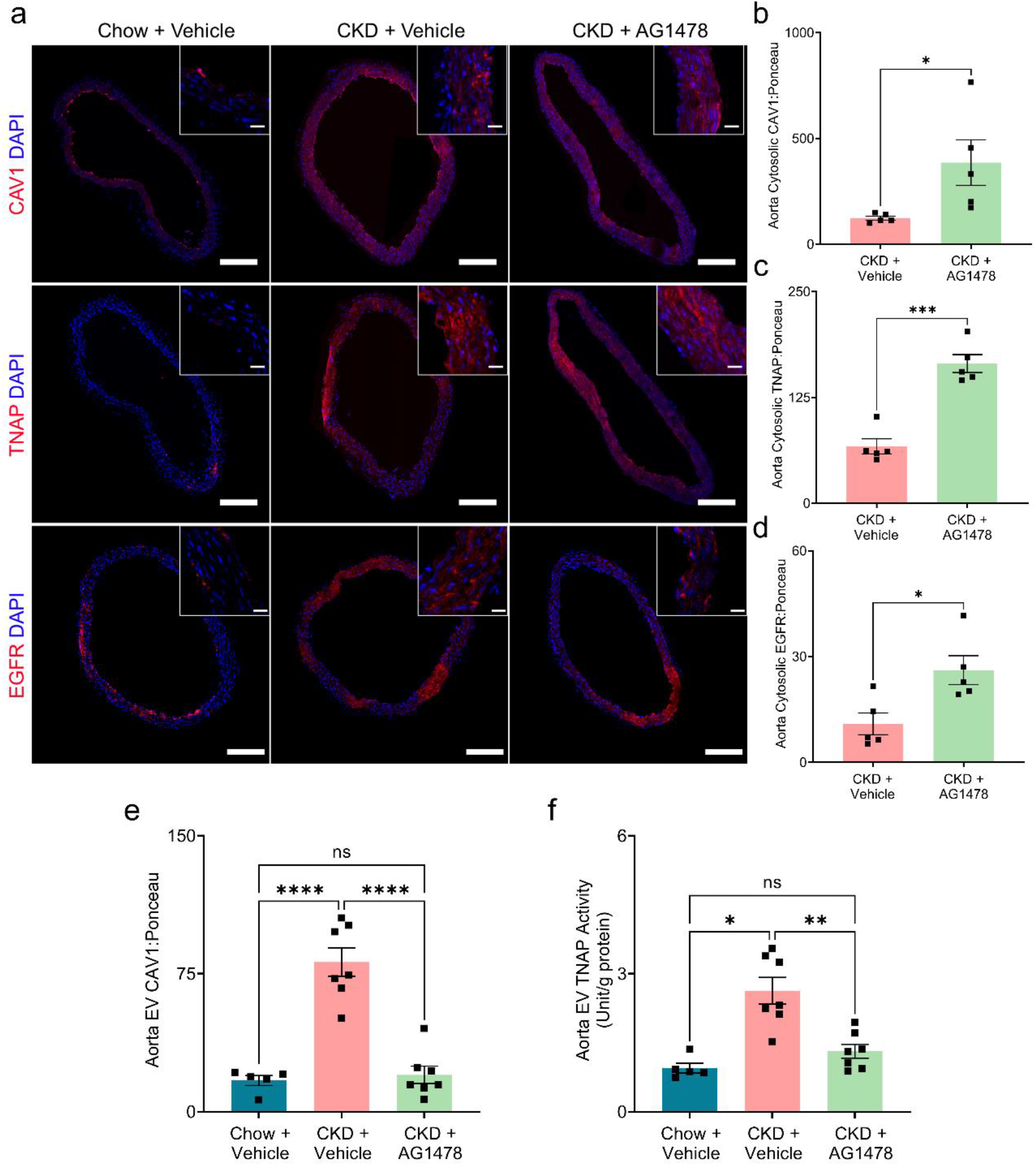
EGFR inhibition redistributes CAV1 and TNAP *in vivo*. (A) Immunofluorescence staining of CAV1 and (B) cytosolic level of CAV1 in aortic tissue; (C) Immunofluorescence staining of TNAP protein and (D) cytosolic level of TNAP protein in aortic tissue; (E) Immunofluorescence staining of EGFR and (F) cytosolic level of EGFR in aortic tissue; EV Level of (G) CAV1 on EVs and (H) TNAP activity isolated from the mouse aortas. (scale bar for 10X and 100X, 200 and 20 μm, respectively). *P < 0.05, **P ≤ 0.01, ***P ≤ 0.001, and ****P ≤ 0.0001, ANOVA with Tukey’s post-hoc test.

### EGFR inhibition reduces the release of CAV1-positive EVs with high TNAP activity in vitro and in vivo

EVs isolated from the aortas of CKD mice exhibited significantly elevated CAV1 protein and TNAP activity compared to chow-fed controls (p < 0.0001 and p = 0.02 for CAV1 and TNAP activity, respectively, **Fig. 3, e** and **f**). The EVs isolated from the CKD mice treated with EGFR inhibitor had significantly lower CAV1 protein and TNAP activity (p < 0.0001 and p = 0.003 for CAV1 and TNAP activity, respectively, **Fig. 3, e** and **f**). The EGFR inhibition led to similar outcomes *in vitro*. EVs obtained from VSMCs cultured in OS media contained significantly elevated CAV1 after 14, 21, and 28 days compared to controls (**Fig. 2**, panel **g**). EV TNAP activity increased in OS VSMC cultures over time (**Fig. 2**, panel **h**). EGFR inhibition significantly reduced the release of EV CAV1 (**Fig. 2**, panel **g**) and EV TNAP activity (**Fig. 2**, panel **h**). Furthermore, EVs isolated from VSMCs cultured in OS media were enriched with Annexin V, a calcium-binding protein, and EGFR (**Fig. 2, i** and **j**); EGFR inhibited groups showed reduced levels of Annexin V and EGFR on the EVs. Of note, the level of CD63, a common exosomal marker, was preserved across the *in vitro* groups following 28 days of culture (p = 0.9 between the groups), as shown in **Fig. 2, k**.

### EGFR inhibition attenuates the interaction between CAV1 and EGFR and redistributes CAV1 in lipid rafts

CAV1 immunoprecipitation showed that in both control and OS cultures, CAV1 interacts with EGFR in VSMCs (**Fig. 4, a**). EGFR inhibition significantly reduced the interaction between CAV1 and EGFR in both control (p = 0.0015) and OS (p < 0.0001) cultures (**Fig. 4, b**). Using ultracentrifugation to perform a density-gradient based separation of VSMC lysates shown that CAV1 redistributes toward more dense fractions in OS cultures with particular enrichment in Fraction 7 compared to control cultures (**Fig. 4, c** and **d**). EGFR inhibition prevented the redistribution of CAV1, leading to a subcellular fractionation profile similar to control cultures (**Fig. 4, d**).

**Figure 4.**
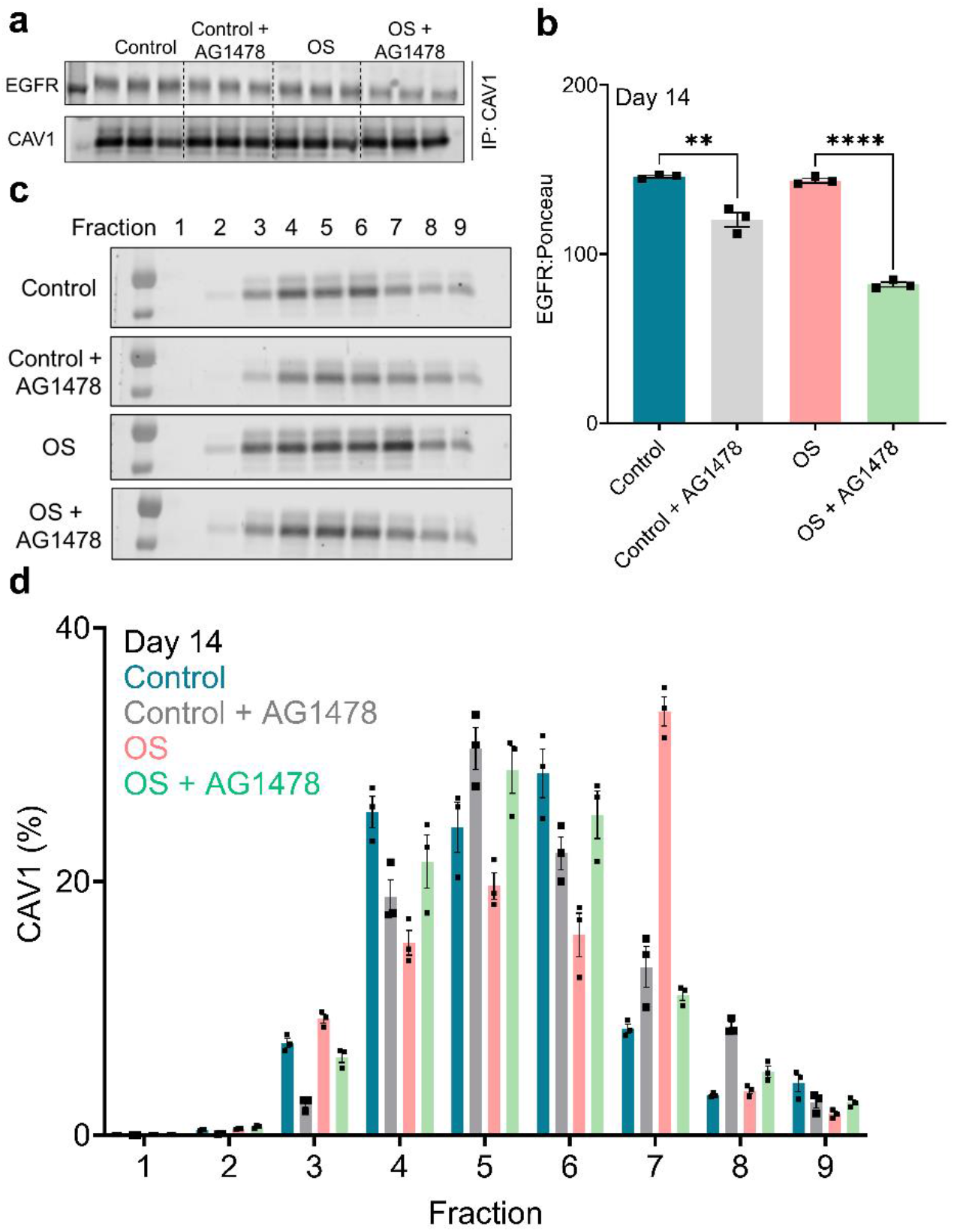
EGFR inhibition attenuates CAV1 and EGFR interaction. (a) EGFR and CAV1 immunoblotting after CAV1 immunoprecipitation from VSMCs following 14 days of treatment; (b) Densitometry and quantification of the EGFR level; (c) CAV1 immunoblotting on isolated lipid rafts; (d) Densitometry and quantification of CAV1 in isolated lipid rafts. \**P* < 0.05, \*\**P* ≤ 0.01, \*\*\**P* ≤ 0.001, and \*\*\*\**P* ≤ 0.0001, ANOVA with Tukey’s post-hoc test.

### EGFR inhibition does not cause deleterious effects on physiological bone mineralization

Both OS and OS cultured osteoblasts treated with EGFR inhibitor committed to osteogenic transition by downregulation of *RUNX2* [34, 35] (**Fig. 5, a**) and increased expression of *ALPL* and *Osteocalcin (BGLAP)* [34], after 7 days (**Fig. 5, b** and **c**), with no significant differences between the groups (p = 0.9 and p = 0.9 for *ALPL* and *BGLAP*, respectively). Similar to *ALPL* expression, the osteoblasts demonstrated significantly increased intracellular TNAP activity after 7 days in both cultures (**Fig. 5, d**). Alizarin red staining demonstrated *in vitro* calcification in both groups and quantification of the *in vitro* calcification showed no significant difference between the groups (p = 0.86, **Fig. 5, e**). In both OS and OS cultured osteoblasts treated with EGFR inhibitor, intracellular CAV1 protein was significantly increased compared to the control group (p = 0.02 and p = 0.01 for the OS and OS with EGFR inhibitor groups, respectively, **Fig. 5, f**). Matrix vesicles released by osteoblasts in both OS and OS treated with EGFR inhibitor groups had significantly increased TNAP activity; however, the EVs from these cells had lower levels of CAV1 protein compared to control on days 14 and 21 in culture (**Fig. 5**, panel **g and h**). We assessed the femurs resected from murine groups to analyze the effects of EGFR inhibition on bone mineralization (**Fig. 6, a** to **c**). The thickness and bone volume fraction of both trabecular (epiphyseal and metaphysical regions) and cortical bone was significantly reduced in CKD animals compared to chow-fed controls. EGFR inhibition increased the thickness of both trabecular and cortical bone significantly in the CKD mice (p = 0.04 and p = 0.02 for epiphyseal and metaphysical regions and p = 0.004 for cortical bone) (**Fig. 6, d** to **f**). Interestingly, EGFR inhibition increased the bone volume fraction in trabecular bone, both epiphyseal (p = 0.009) and metaphysical (p = 0.002) regions, compared to CKD animals. However, it did not significantly change in cortical bone (p = 0.25) (**Fig. 6, g** to **i**). Detailed quantification of the bone structural parameters can be found in **Online materials, Table I**.

**Figure 5.**
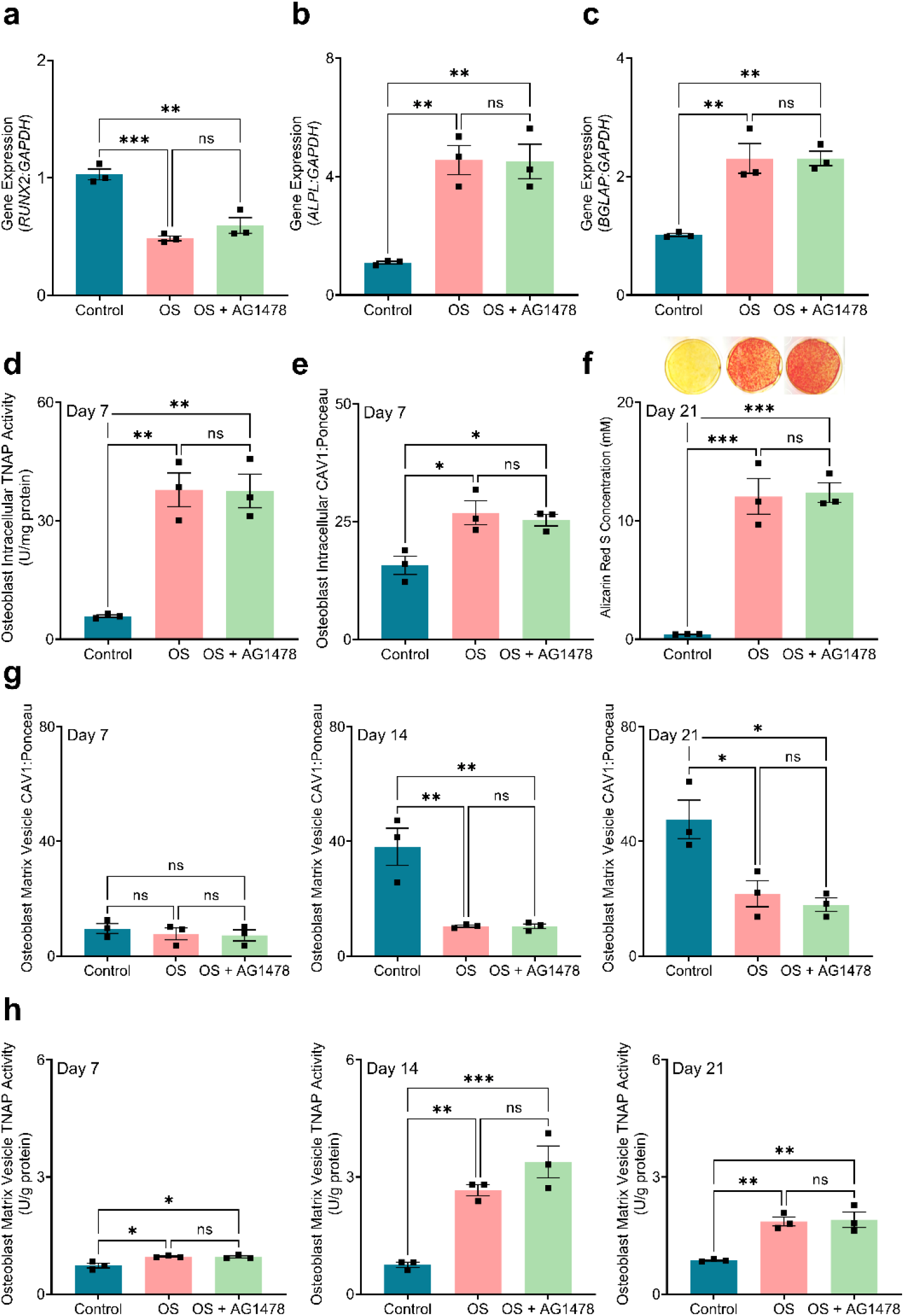
EGFR inhibition does not prevent osteoblast *in vitro* calcification. (a, b, and c) Gene expression of common osteogenic markers, *RUNX2, ALPL*, and *BGLAP* in osteoblasts following 7 days of treatment; (d) Osteoblast intracellular TNAP activity following 7 days of treatment; (e) Alizarin Red S staining and quantification of osteoblast cultures after 21 days; (f) Osteoblast intracellular CAV1 following 7 days of treatment; (g) CAV1 level on matrix vesicles liberated from osteoblasts on days 7, 14, and 21 of culture; (h) TNAP activity of matrix vesicles isolated from osteoblast cultures on days 7, 14, and 21. \**P* < 0.05, \*\**P* ≤ 0.01, \*\*\**P* ≤ 0.001, and \*\*\*\**P* ≤ 0.0001, ANOVA with Tukey’s post-hoc test.

**Figure 6.**
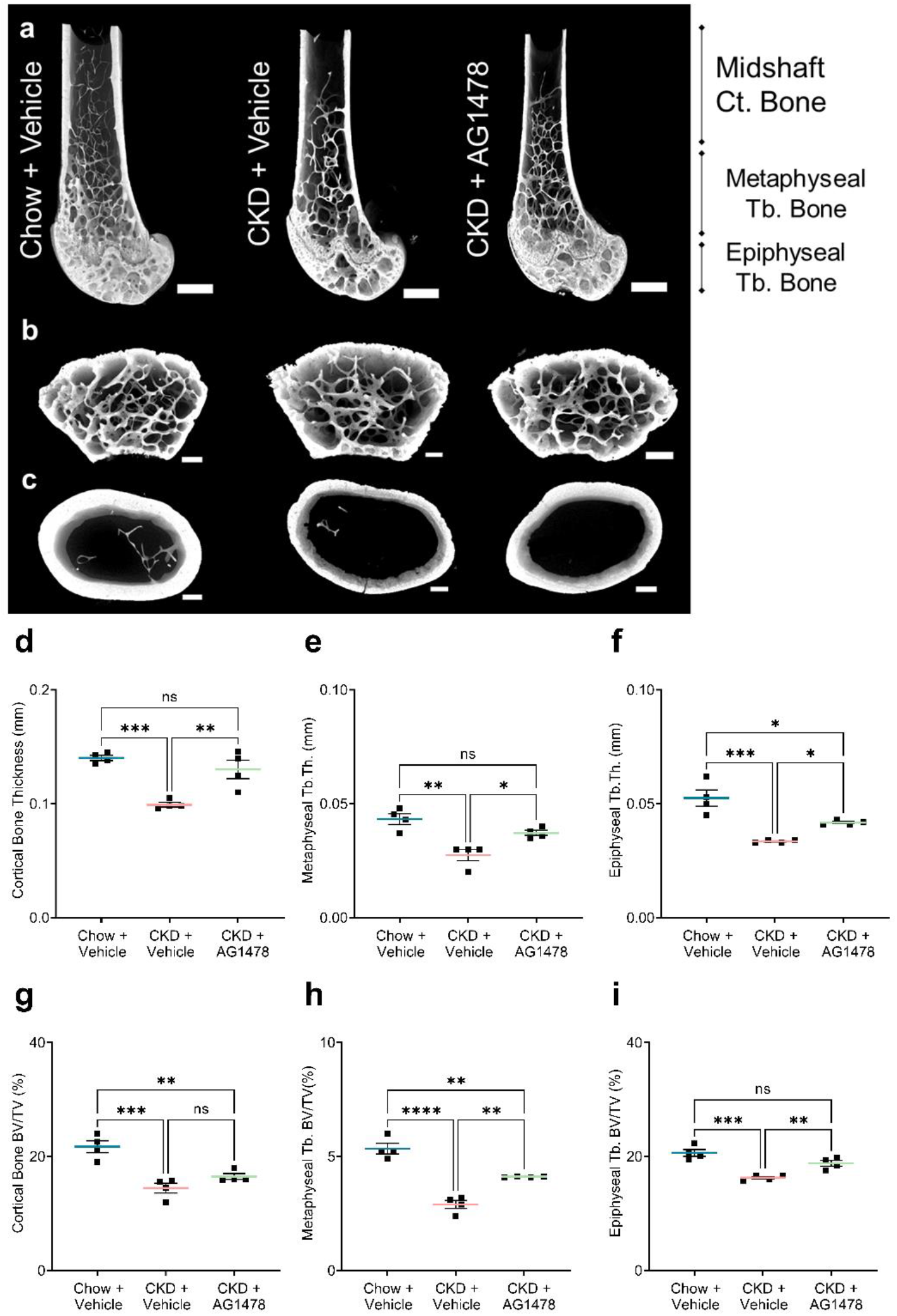
EGFR inhibition does not have deleterious effects on physiological bone mineralization. 3D reconstructions of (a) femoral head, (b) cancellous bone, and (c) cortical bone resected from mouse groups (scale bar: 0.5 mm); Bone thickness at: (d) Cortical, (e) Metaphyseal trabecular, and (f) Epiphyseal trabecular regions; Bone volume fraction (%) at: (g) Cortical, (h) Metaphyseal trabecular, and (i) Epiphyseal trabecular regions. \**P* < 0.05, \*\**P* ≤ 0.01, \*\*\**P* ≤ 0.001, and \*\*\*\**P* ≤ 0.0001, ANOVA with Tukey’s post-hoc test.

### Mendelian Randomization shows positive correlation between serum EGFR and CAC

Of the 11 MR regressions performed in the MESA cohort, all regressions predicted positive correlation between serum EGFR concentration and CAC (i.e., elevated EGFR concentration predicts increased incidence of elevated CAC). Two of the MR regressions reached statistical significance beyond the Bonferroni-adjusted significance threshold: robust MR-Egger and penalized robust MR-Egger. The intercept tests for the MR-Egger estimates are statistically significant at p = 1.9×10^-5^, suggesting the presence of vertical pleiotropy among the IV SNPs accounted for in the MR-Egger type regressions (**Fig. 7, a**). The causal estimates of the effect of EGFR concentration on increased CAC are associated with p-values < 1×10^-10^ (**Online materials**, **Table II**).

**Figure 7.**
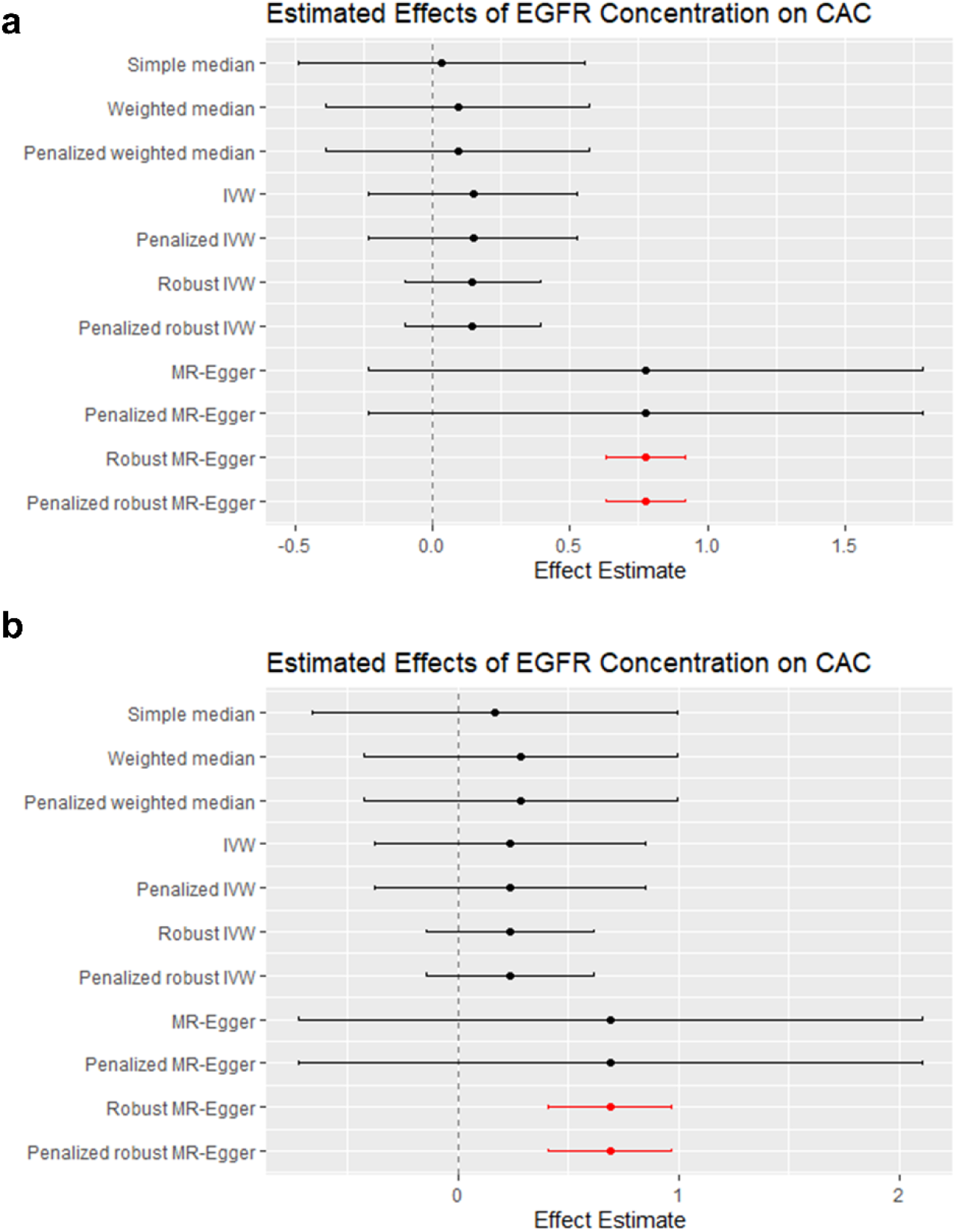
Clinical data indicate positive correlation between serum EGFR and coronary artery calcification. Forest plot summarizing effect estimates of each MR regression along with their 95% confidence intervals for: (a) MESA cohort; (b) Offspring cohort of FHS. Robust MR-Egger and penalized robust MR-Egger estimates of effect are statistically significant and highlighted in red.

Replication of the 11 MR regressions in the FHS cohort also yielded significant estimates for the robust MR-Egger and penalized robust MR-Egger regression estimates with p-values of 1.17×10^-6^ for both (**Fig. 7, b**). However, the intercept test for vertical pleiotropy was not statistically significant (p = 0.06), possibly trending towards significance due to insufficient sample size. However, both regressions suggest a positive causal relation between serum EGFR concentration and CAC (**Online materials, Table III**).

## Discussion

Despite the recognition that EVs participate in vascular calcification, significant knowledge gaps exist into how these specialized structures form and to what extent they mediate mineral deposition. Here, we present new insight into the role of CAV1 trafficking in the formation of calcifying EVs and demonstrate that EGFR inhibition can alter CAV1 trafficking to prevent vascular calcification. Caveolae have a low buoyant density. Translocation of caveolin-1 to more dense regions in the gradient-based VSMC fractionation analyses indicates trafficking to non-caveolar domains [36] in pro-calcific conditions (**Fig. 4, d**). The data presented in the current study suggest that physical interactions between EGFR and caveolin-1 are required for the intracellular, non-caveolar trafficking mechanisms that lead to calcifying EV biogenesis. EGFR tyrosine kinase inhibition reduces EGFR-CAV1 co-immunoprecipitation (**Fig. 4, b**) and retains CAV1 within less dense membrane fractions associated with caveolae (**Fig. 4, d**). The EGFR inhibition, however, does not alter transition of VSMCs to a pro-calcifying phenotype (**Fig. 1, k** to **m**). By preventing the formation of calcifying EVs, factors expressed during this phenotypic transition accumulate within the VSMCs (**Fig. 2, e** to **f** and **3, b to d**). Blocking the release of the pro-calcific factors in EVs resulted in reduced calcification *in vitro* (**Fig. 1, i**) and *in vivo* (**Fig. 1, a**), demonstrating the relevance of calcifying EVs in mineral formation.

Given the myriad of upstream initiators of vascular calcification, altering calcifying EV biogenesis may represent a point of convergence that can be targeted therapeutically [37]. CKD patients are particularly prone to develop widespread vascular calcification, increasing from 25% of patients in stages 3 and 4 to 50-80% of the population in stage 5 [38]. Our data indicate that CKD-induced vascular calcification associates with increased serum EGFR in mice, similar to the association between circulating EGFR and CAC in the MR regression analyses. Osteogenic transition of VSMCs in the vessel wall and release of calcifying EVs orchestrate progression of vascular calcification [38]. These EVs share similarities with matrix vesicles released by osteoblasts and chondrocytes [3]. The plasma membrane protein, CAV1, plays a key role in formation of calcifying EVs and VSMC-mediated calcification. Knockdown of CAV1 led to prevention of VSMCs calcification *in vitro* [11]. EGFR co-localizes and interacts with CAV1 in caveolar domains of the plasma membrane. Previous studies showed that EGFR facilitates tyrosine kinase mediated phosphorylation of CAV1 and modulates CAV1 trafficking [39–42]. Therefore, we hypothesized that EGFR tyrosine kinase inhibition may prevent the CAV1-dependent formation of calcifying EVs in CKD.

Here, we show that inhibiting EGFR tyrosine kinase activity prevents vascular calcification in a CKD mouse model, with 100% survival rate. The *in vivo* results showed reduced calcium burden in the aorta of CKD mice treated with EGFR tyrosine kinase inhibitor, AG1478. This effect was independent of kidney remodeling as AG1478 treatment did not reduce the expression of common markers of renal injury. Furthermore, previous reports indicated the correlation between elevated serum TNAP activity and renal injury due to endothelial dysfunction and inflammation [43, 44]. High serum TNAP activity correlates with vascular calcification and dysfunctional bone turnover in CKD [45]. Indeed, TNAP facilitates the hydrolysis of a common calcification inhibitor, inorganic pyrophosphate, and mediates the calcification process [43, 46]. We showed similar elevated serum TNAP activity in both CKD mice and CKD mice treated with EGFR inhibitor, demonstrating that EGFR inhibition prevented vascular calcification independent from serum TNAP activity and renal injury.

We hypothesized that EGFR inhibition would prevent calcification by altering CAV1 trafficking and disrupting the biogenesis of calcifying EVs. Our findings demonstrated elevated CAV1-positive EVs in the aorta of CKD mice, which was reduced by EGFR inhibition. Similarly, TNAP activity was elevated in EVs isolated from the aortae of CKD mice, while EGFR inhibition reduced the activity of this enzyme in the EVs. Calcifying EVs are enriched in Annexin V, a collagen-binding Ca^2+^ channel [3, 47]. We found that Annexin V was elevated in VSMC EVs, which was also reduced by EGFR inhibition. Taken together, these results support our hypothesis and suggest that targeting the CAV1-dependent formation of calcifying EVs by EGFR inhibition reduced vascular calcification in the CKD mouse model.

Since our data indicate that EGFR inhibition disrupts calcifying EV formation, we also set out to determine whether the treatment alters other types of EV formation. We began by blotting for CD63, a widely utilized marker enriched in exosomes and other EV subtypes, including high phosphate-induced VSMC calcification [48]. The data demonstrate no differences in CD63 protein within EVs from VSMCs cultured in control, OS, or OS media samples treated with AG1478. It is unclear whether calcifying EVs considered in our study derive from a CD63-positive population that is loaded with pro-calcific components, or whether they derive from a distinct population of EVs. Though these data do not show changes in CD63, it is possible that CD63-positive vesicles acquire pro-calcific properties in pathological conditions. The current data, however, suggest that CD63-positive EV release is not altered by EGFR inhibition.

Our data also suggest that osteogenic function of osteoblasts was not affected by EGFR inhibition. Culturing osteoblasts in OS media resulted in the release of TNAP-positive EVs and robust mineralization, neither of which was altered by EGFR inhibition. Interestingly, we showed reduced CAV1 levels in matrix vesicles released by osteoblasts cultured in OS media. These observations further suggest that, despite many commonalities, bone matrix vesicles and vascular calcifying EVs originate through different mechanisms. CKD patients often exhibit bone disorders, including decreased bone mass density [49]. Previous reports demonstrated that trabecular and cortical bone mass density increased in CAV1-deficient mice [50, 51]. Here, we demonstrated that EGFR inhibition significantly reversed reductions in trabecular and cortical thickness in the CKD mice; bone volume fraction in trabecular regions significantly increased by the treatment, while cortical bone volume fraction was not improved. At the least, these results suggest that EGFR inhibition does not induce deleterious bone remodeling and—at best—may improve CKD-induced bone pathologies. The calcification paradox—the observation that bone and vascular mineral are often negatively correlated [52]—is poorly understood. Future studies that further explore the role of CAV1 and EGFR in calcification may provide new mechanistic insight into physiological and pathological mineralization differences.

We also assessed the relevance of our findings with clinical outcomes using cardioinformatics techniques. Mendelian Randomization is a causal inference technique used for the *in silico* identification of novel drug targets, potential drug-drug interactions/synergies, as well as for estimation of magnitudes of effect for each of these [53]. MR uses genetic variants as IVs, where the unique genetic composition of each individual is used to “randomize” individuals into different treatment groups, mimicking a randomized control trial since genetic composition is randomized at birth [54]. The intuition in MR is that if genetic variants are correlated with the exposure variable and if the exposure variable is causal for the outcome variable, then the genetic variants should also explain variance in the outcome variable [55]. The IV assumptions must be fulfilled for MR to yield valid results, though in practice, they are often violated due to pleiotropic effects of genetic variants [30]. Therefore, a wide set of MR regression techniques have been developed, each with unique merits in accounting for minimizing and resisting potential violations of IV assumptions or other confounding factors [30]. As the number of tools created to support *in silico* target discovery continues increasing, in particular database tools such as HeartBioPortal, OpenGWAS, and MRBase, MR becomes an increasingly attractive tool for exploration of novel pharmaceutical interventions [56–59].

The direction of effect was qualitatively replicated in each of the MR regressions in the FHS cohort. The significant causal estimates of the robust MR-Egger and penalized robust MR-Egger estimates were recreated, though the intercept test p-value for the regressions fell short of reaching statistical significance in the FHS cohort. The intercept p-values for the two MR-Egger regressions with significant causal estimates were p = 0.06, which likely did not reach statistical significance due to lower sample size of our FHS replication cohort (n = 1,896) relative to our MESA discovery cohort (n = 5,755). We interpreted the highly significant positive causal estimates and intercepts in the MESA cohort (p < 1 x10^-10^ and p < 1.9 x10^-5^) along with the significant positive causal estimates and non-significant intercept (p = 1.17×10^-6^ and p = 0.06) in the FHS cohort as suggestive that increased serum EGFR is causal for increased CAC. This cardioinformatics workflow [60] highlights the importance of bridging not only the bench-to-bedside but also the informatics-to-medicine divide that still exists in modern precision cardiology research. This approach can connect basic science to population-level data and enable computationally-derived therapeutics.

## Conclusion

Cardiovascular disease is the leading cause of death in patients with CKD, and the risk of mortality is directly associated with the presence of vascular calcification. Therefore, the development of a therapeutic strategy to prevent vascular mineralization in these patients would represent a breakthrough in CKD management. Other therapeutic strategies in promising clinical trials slow CKD-mediated vascular calcification by interacting directly with mineral [61]. Other proposed pre-clinical strategies include targeting mechanisms that lead to a pro-calcific SMC phenotype. However, a myriad of initiators results in vascular calcification. Our data suggest that EGFR inhibition does not alter SMC phenotype, but directly affects caveolin-1 trafficking. This provides a unique therapeutic strategy to modulate calcifying EV formation independent of cell phenotype. EGFR inhibitors have demonstrated clinical safety and efficacy in cancer treatments [62]. The accessibility of EGFR has led to the suggestion that it may represent a therapeutic target worth exploring for cardiovascular diseases [63]. CKD patients represent an identifiable population in need of therapeutics for vascular calcification. The confluence of an accessible target with approved therapeutics and a clear patient population that lack therapeutic options could accelerate the start of clinical trials.

## Acknowledgements

MESA data is accessed under Database of Genotypes and Phenotypes (dbGaP) accession phs000209.v13.p3. MESA and the MESA SHARe project are conducted and supported by the National Heart, Lung, and Blood Institute (NHLBI) in collaboration with MESA investigators. Support for MESA is provided by contracts N01-HC95159, N01-HC-95160, N01-HC-95161, N01-HC-95162, N01-HC-95163, N01-HC-95164, N01-HC-95165, N01-HC95166, N01-HC-95167, N01-HC-95168, N01-HC-95169, UL1-RR-025005, and UL1-TR-000040. Funding for SHARe genotyping was provided by NHLBI Contract N02-HL-64278. Genotyping was performed at Affymetrix (Santa Clara, California, USA) and the Broad Institute of Harvard and MIT (Boston, Massachusetts, USA) using the Affymetrix Genome-Wide Human SNP Array 6.0. This work was completed in part with resources provided by the University of Chicago Research Computing Center.

## References

1. Ho, C.Y. and C.M. Shanahan, Medial arterial calcification: an overlooked player in peripheral arterial disease. Arterioscler. Thromb. Vasc. Biol., 2016. 36(8): p. 1475–1482.

2. Marinelli, A., V. Pistolesi, L. Pasquale, L. Di Lullo, M. Ferrazzano, G. Baudena, F. Della Grotta, and A. Di Napoli, Diagnosis of arterial media calcification in chronic kidney disease. Cardiorenal Med., 2013. 3(2): p. 89–95.

3. Bakhshian Nik, A., J.D. Hutcheson, and E. Aikawa, Extracellular vesicles as mediators of cardiovascular calcification. Front. cardiovasc. med., 2017. 4: p. (78)1–12.

4. Manzoor, S., S. Ahmed, A. Ali, K.H. Han, I. Sechopoulos, A. O’Neill, B. Fei, and W.C. O’Neill, Progression of medial arterial calcification in CKD. Kidney Int. Rep., 2018. 3(6): p. 1328–1335.

5. London, G.M., A.P. Guerin, S.J. Marchais, F. Metivier, B. Pannier, and H. Adda, Arterial media calcification in end-stage renal disease: impact on all-cause and cardiovascular mortality. Nephrol. Dial. Transplant., 2003. 18(9): p. 1731–40.

6. Moe, S.M. and N.X. Chen, Mechanisms of vascular calcification in chronic kidney disease. J. Am. Soc. Nephrol., 2008. 19(2): p. 213–216.

7. Ruiz, J.L., J.D. Hutcheson, and E. Aikawa, Cardiovascular calcification: current controversies and novel concepts. Cardiovasc. Pathol., 2015. 24(4): p. 207–212.

8. New, S.E. and E. Aikawa, Role of extracellular vesicles in de novo mineralization: an additional novel mechanism of cardiovascular calcification. Arterioscler. Thromb. Vasc. Biol., 2013. 33(8): p. 1753–1758.

9. Shapiro, I.M., W.J. Landis, and M.V. Risbud, Matrix vesicles: Are they anchored exosomes? Bone, 2015. 79: p. 29–36.

10. Aikawa, E. and J.D. Hutcheson, Cardiovascular Calcification and Bone Mineralization. 2020: Springer.

11. Goettsch, C., J.D. Hutcheson, M. Aikawa, H. Iwata, T. Pham, A. Nykjaer, M. Kjolby, M. Rogers, T. Michel, and M. Shibasaki, Sortilin mediates vascular calcification via its recruitment into extracellular vesicles. J. Clin. Investig., 2016. 126(4): p. 1323–1336.

12. Hardin, C.D. and J. Vallejo, Caveolins in vascular smooth muscle: form organizing function. Cardiovasc. Res., 2006. 69(4): p. 808–815.

13. Gratton, J.-P., P. Bernatchez, and W.C. Sessa, Caveolae and caveolins in the cardiovascular system. Circ. Res., 2004. 94(11): p. 1408–1417.

14. Liu, P., M. Rudick, and R.G. Anderson, Multiple functions of caveolin-1. J. Biol. Chem, 2002. 277(44): p. 41295–41298.

15. Wieduwilt, M. and M. Moasser, The epidermal growth factor receptor family: biology driving targeted therapeutics. Cell. Mol. Life Sci., 2008. 65(10): p. 1566–1584.

16. Zhang, Y., F. Peng, B. Gao, A.J. Ingram, and J.C. Krepinsky, Mechanical strain-induced RhoA activation requires NADPH oxidase-mediated ROS generation in caveolae. Antioxid. Redox Signal, 2010. 13(7): p. 959–973.

17. Zhang, B., F. Peng, D. Wu, A.J. Ingram, B. Gao, and J.C. Krepinsky, Caveolin-1 phosphorylation is required for stretch-induced EGFR and Akt activation in mesangial cells. Cell. Signal., 2007. 19(8): p. 1690–1700.

18. Wykosky, J., T. Fenton, F. Furnari, and W.K. Cavenee, Therapeutic targeting of epidermal growth factor receptor in human cancer: successes and limitations. Chin. J. Cancer, 2011. 30(1): p. 5.

19. Liang, Y.N., Y. Liu, L. Wang, G. Yao, X. Li, X. Meng, F. Wang, M. Li, D. Tong, and J. Geng, Combined caveolin-1 and epidermal growth factor receptor expression as a prognostic marker for breast cancer. Oncol. Lett., 2018. 15(6): p. 9271–9282.

20. Shin, S.U., J. Lee, J.H. Kim, W.H. Kim, S.E. Song, A. Chu, H.S. Kim, W. Han, H.S. Ryu, and W.K. Moon, Gene expression profiling of calcifications in breast cancer. Sci. Rep., 2017. 7(1): p. 1–11.

21. Wang, L., Z. Huang, W. Huang, X. Chen, P. Shan, P. Zhong, Z. Khan, J. Wang, Q. Fang, and G. Liang, Inhibition of epidermal growth factor receptor attenuates atherosclerosis via decreasing inflammation and oxidative stress. Sci. Rep., 2017. 7(1): p. 1–14.

22. Tani, T., H. Orimo, A. Shimizu, and S. Tsuruoka, Development of a novel chronic kidney disease mouse model to evaluate the progression of hyperphosphatemia and associated mineral bone disease. Sci. Rep., 2017. 7(1): p. 1–12.

23. Chen, N.X., K. O’neill, X. Chen, K. Kiattisunthorn, V.H. Gattone, and S.M. Moe, Transglutaminase 2 accelerates vascular calcification in chronic kidney disease. Am. J. Nephrol, 2013. 37(3): p. 191–198.

24. Goettsch, C., M. Rauner, N. Pacyna, U. Hempel, S.R. Bornstein, and L.C. Hofbauer, miR-125b regulates calcification of vascular smooth muscle cells. Am. J. Pathol., 2011. 179(4): p. 1594–1600.

25. Hutcheson, J.D., C. Goettsch, S. Bertazzo, N. Maldonado, J.L. Ruiz, W. Goh, K. Yabusaki, T. Faits, C. Bouten, and G. Franck, Genesis and growth of extracellular-vesicle-derived microcalcification in atherosclerotic plaques. Nature materials, 2016. 15(3): p. 335–343.

26. Ng, H.H., M. Jelinic, L.J. Parry, and C.-H. Leo, Increased superoxide production and altered nitric oxide-mediated relaxation in the aorta of young but not old male relaxin-deficient mice. AM J PHYSIOL-HEART C, 2015. 309(2): p. H285–H296.

27. Ng, H.H., C.H. Leo, and L.J. Parry, Serelaxin (recombinant human relaxin-2) prevents high glucose-induced endothelial dysfunction by ameliorating prostacyclin production in the mouse aorta. Pharmacol. Res., 2016. 107: p. 220–228.

28. Ng, H.H., C.H. Leo, D. Prakoso, C. Qin, R.H. Ritchie, and L.J. Parry, Serelaxin treatment reverses vascular dysfunction and left ventricular hypertrophy in a mouse model of Type 1 diabetes. Sci. Rep., 2017. 7(1): p. 1–15.

29. Steiner, L., A. Synek, and D.H. Pahr, Comparison of different microCT-based morphology assessment tools using human trabecular bone. Bone reports, 2020. 12: p. 100261.

30. Burgess, S., G.D. Smith, N.M. Davies, F. Dudbridge, D. Gill, M.M. Glymour, F.P. Hartwig, M.V. Holmes, C. Minelli, and C.L. Relton, Guidelines for performing Mendelian randomization investigations. Wellcome Open Research, 2019. 4.

31. Sun, B.B., J.C. Maranville, J.E. Peters, D. Stacey, J.R. Staley, J. Blackshaw, S. Burgess, T. Jiang, E. Paige, and P. Surendran, Genomic atlas of the human plasma proteome. Nature, 2018. 558(7708): p. 73–79.

32. Yavorska, O.O. and S. Burgess, MendelianRandomization: an R package for performing Mendelian randomization analyses using summarized data. International journal of epidemiology, 2017. 46(6): p. 1734–1739.

33. RCoreTeam, R: A language and environment for statistical computing. 2021, R Foundation for Statistical Computing, Vienna, Austria.

34. Rutkovskiy, A., K.-O. Stensløkken, and I.J. Vaage, Osteoblast differentiation at a glance. Med. Sci. Monit. Basic Res., 2016. 22: p. 95.

35. Komori, T., Regulation of bone development and extracellular matrix protein genes by RUNX2. Cell and tissue research, 2010. 339(1): p. 189–195.

36. Kawabe, J., S. Okumura, M.C. Lee, J. Sadoshima, and Y. Ishikawa, Translocation of caveolin regulates stretch-induced ERK activity in vascular smooth muscle cells. Am J Physiol Heart Circ Physiol, 2004. 286(5): p. H1845–52.

37. Ruiz, J.L., S. Weinbaum, E. Aikawa, and J.D. Hutcheson, Zooming in on the genesis of atherosclerotic plaque microcalcifications. J Physiol, 2016. 594(11): p. 2915–27.

38. Dusso, A., M.I. Colombo, and C.M. Shanahan, Not all vascular smooth muscle cell exosomes calcify equally in chronic kidney disease. Kidney Int., 2018. 93(2): p. 298–301.

39. Kim, Y.-N., G.J. Wiepz, A.G. Guadarrama, and P.J. Bertics, Epidermal growth factor-stimulated tyrosine phosphorylation of caveolin-1: enhanced caveolin-1 tyrosine phosphorylation following aberrant epidermal growth factor receptor status. J. Biol. Chem, 2000. 275(11): p. 7481–7491.

40. Abulrob, A., S. Giuseppin, M.F. Andrade, A. McDermid, M. Moreno, and D. Stanimirovic, Interactions of EGFR and caveolin-1 in human glioblastoma cells: evidence that tyrosine phosphorylation regulates EGFR association with caveolae. Oncogene, 2004. 23(41): p. 6967–6979.

41. Dittmann, K., C. Mayer, R. Kehlbach, and H.P. Rodemann, Radiation-induced caveolin-1 associated EGFR internalization is linked with nuclear EGFR transport and activation of DNA-PK. Molecular cancer, 2008. 7(1): p. 1–9.

42. Wang, Y., O. Roche, C. Xu, E.H. Moriyama, P. Heir, J. Chung, F.C. Roos, Y. Chen, G. Finak, and M. Milosevic, Hypoxia promotes ligand-independent EGF receptor signaling via hypoxia-inducible factor-mediated upregulation of caveolin-1. Proceedings of the National Academy of Sciences, 2012. 109(13): p. 4892–4897.

43. Sciacqua, A., G. Tripepi, M. Perticone, V. Cassano, T.V. Fiorentino, G.N. Pititto, R. Maio, S. Miceli, F. Andreozzi, and G. Sesti, Alkaline phosphatase affects renal function in never-treated hypertensive patients: effect modification by age. Sci. Rep., 2020. 10(1): p. 1–7.

44. Perticone, F., M. Perticone, R. Maio, A. Sciacqua, M. Andreucci, G. Tripepi, S. Corrao, F. Mallamaci, G. Sesti, and C. Zoccali, Serum alkaline phosphatase negatively affects endothelium-dependent vasodilation in naive hypertensive patients. Hypertension, 2015. 66(4): p. 874–880.

45. Taliercio, J.J., J.D. Schold, J.F. Simon, S. Arrigain, A. Tang, G. Saab, J.V. Nally Jr, and S.D. Navaneethan, Prognostic importance of serum alkaline phosphatase in CKD stages 3-4 in a clinical population. American journal of kidney diseases, 2013. 62(4): p. 703–710.

46. Schoppet, M. and C. Shanahan, Role for alkaline phosphatase as an inducer of vascular calcification in renal failure? Kidney Int., 2008. 73(9): p. 989–991.

47. Chen, N.X., K.D. O’Neill, X. Chen, and S.M. Moe, Annexin-mediated matrix vesicle calcification in vascular smooth muscle cells. Journal of Bone and Mineral Research, 2008. 23(11): p. 1798–1805.

48. Kapustin, A.N., M.L. Chatrou, I. Drozdov, Y. Zheng, S.M. Davidson, D. Soong, M. Furmanik, P. Sanchis, R.T.M. De Rosales, and D. Alvarez-Hernandez, Vascular smooth muscle cell calcification is mediated by regulated exosome secretion. Circ. Res., 2015. 116(8): p. 1312–1323.

49. Pan, B.-L. and S.-S. Loke, Chronic kidney disease associated with decreased bone mineral density, uric acid and metabolic syndrome. PloS one, 2018. 13(1): p. e0190985.

50. Rubin, J., Z. Schwartz, B.D. Boyan, X. Fan, N. Case, B. Sen, M. Drab, D. Smith, M. Aleman, and K.L. Wong, Caveolin-1 knockout mice have increased bone size and stiffness. Journal of Bone and Mineral Research, 2007. 22(9): p. 1408–1418.

51. Lee, Y.D., S.-H. Yoon, C.K. Park, J. Lee, Z.H. Lee, and H.-H. Kim, Caveolin-1 regulates osteoclastogenesis and bone metabolism in a sex-dependent manner. J. Biol. Chem, 2015. 290(10): p. 6522–6530.

52. Persy, V. and P. D’Haese, Vascular calcification and bone disease: the calcification paradox. Trends. Mol. Med., 2009. 15(9): p. 405–416.

53. Gill, D., M.K. Georgakis, V.M. Walker, A.F. Schmidt, A. Gkatzionis, D.F. Freitag, C. Finan, A.D. Hingorani, J.M. Howson, and S. Burgess, Mendelian randomization for studying the effects of perturbing drug targets. Wellcome Open Research, 2021. 6.

54. Lawlor, D.A., R.M. Harbord, J.A. Sterne, N. Timpson, and G. Davey Smith, Mendelian randomization: using genes as instruments for making causal inferences in epidemiology. Statistics in medicine, 2008. 27(8): p. 1133–1163.

55. Sun, S., Y. Liu, L. Li, M. Jiao, Y. Jiang, B. Li, W. Gao, and X. Li, Mendelian randomization analysis of the association between human blood cell traits and uterine polyps. Scientific reports, 2021. 11(1): p. 1–9.

56. Khomtchouk, B.B., K.A. Vand, W.C. Koehler, D.-T. Tran, K. Middlebrook, S. Sudhakaran, C.S. Nelson, O. Gozani, and T.L. Assimes, HeartBioPortal: an internet-of-omics for human cardiovascular disease data. Circulation: Genomic and Precision Medicine, 2019. 12(4): p. e002426.

57. Khomtchouk, B.B., C.S. Nelson, K.A. Vand, S. Palmisano, and R.L. Grossman, HeartBioPortal2. 0: new developments and updates for genetic ancestry and cardiometabolic quantitative traits in diverse human populations. Database, 2020. 2020.

58. Hemani, G., J. Zheng, B. Elsworth, K.H. Wade, V. Haberland, D. Baird, C. Laurin, S. Burgess, J. Bowden, and R. Langdon, The MR-Base platform supports systematic causal inference across the human phenome. elife, 2018. 7: p. e34408.

59. Elsworth, B., M. Lyon, T. Alexander, Y. Liu, P. Matthews, J. Hallett, P. Bates, T. Palmer, V. Haberland, G.D. Smith, J. Zheng, P. Haycock, T. Gaunt, and G. Hemani, The MRC IEU OpenGWAS data infrastructure. 2020, bioRxiv.

60. Khomtchouk, B.B., D.-T. Tran, K.A. Vand, M. Might, O. Gozani, and T.L. Assimes, Cardioinformatics: the nexus of bioinformatics and precision cardiology. Briefings in bioinformatics, 2020. 21(6): p. 2031–2051.

61. Raggi, P., A. Bellasi, D. Bushinsky, J. Bover, M. Rodriguez, M. Ketteler, S. Sinha, C. Salcedo, K. Gillotti, C. Padgett, R. Garg, A. Gold, J. Perello, and G.M. Chertow, Slowing Progression of Cardiovascular Calcification With SNF472 in Patients on Hemodialysis: Results of a Randomized Phase 2b Study. Circulation, 2020. 141(9): p. 728–739.

62. Seshacharyulu, P., M.P. Ponnusamy, D. Haridas, M. Jain, A.K. Ganti, and S.K. Batra, Targeting the EGFR signaling pathway in cancer therapy. Expert Opin. Ther. Targets, 2012. 16(1): p. 15–31.

63. Mindur, J.E. and F.K. Swirski, Growth factors as immunotherapeutic targets in cardiovascular disease. Arterioscler. Thromb. Vasc. Biol., 2019. 39(7): p. 1275–1287.

