## Supplemental Material for "Epidermal Growth Factor Receptor Inhibition Prevents Caveolin-1-dependent Calcifying Extracellular Vesicle Biogenesis"

Or

Bohdan B. Khomtchouk, PhD
900 East 57th Street, KCBD 10150
Chicago, IL 60637, USA.


**This file includes**:
Supplementary Table I to III
Supplementary Figure I and III

**Supplementary Table I**. Detailed bone structural parameters.

| Parameter | Chow + Vehicle n = 4 | CKD + Vehicle n = 4 | CKD + AG1478 n = 4 |
|---|---|---|---|
| **Distal femur** | | | |
| *Epiphyseal Trabecular Bone* | | | |
| BV/TV (%) | 20.64±0.5 | 16.275±0.192 | 18.825±0.425 |
| Tb.N ($mm^{-1}$) | 5.16±0.11 | 4.92±0.25 | 5±0.19685 |
| Tb.Th (mm) | 0.052±0.003 | 0.0335±0.0002 | 0.0417±0.0004 |
| Tb.Sp (mm) | 0.417±0.007 | 0.468±0.0106 | 0.450±0.0132 |
| BS/BV ($mm^2/mm^3$) | 65.32±5.52 | 87.77±11.896 | 81.82±5.7973 |
| EF | 0.0215±0.0009 | 0.0252±0.0012 | 0.0227±0.0006 |
| $EF_{max}$ | 0.876±0.007 | 0.8437±0.0082 | 0.8513±0.007 |
| $EF_{min}$ | -0.805±0.008 | -0.756±0.0175 | -0.776±0.02 |
| DA | 1.462±0.030 | 1.263±0.0173 | 1.351±0.02 |
| *Metaphyseal Trabecular Bone* | | | |
| BV/TV (%) | 5.35±0.2 | 2.9±0.15 | 4.1±0.008 |
| Tb.N ($mm^{-1}$) | 4.92±0.19 | 4.08±0.13 | 4.39±0.12 |
| Tb.Th (mm) | 0.043±0.002 | 0.027±0.002 | 0.037±0.001 |
| Tb.Sp (mm) | 0.408±0.02 | 0.472±0.013 | 0.44±0.017 |
| BS/BV ($mm^2/mm^3$) | 79.65±1.86 | 94.19±3.2 | 90.65±3.37 |
| EF | 0.086±0.004 | 0.117±0.006 | 0.098±0.004 |
| $EF_{max}$ | 0.86±0.008 | 0.73±0.05 | 0.81±0.004 |
| $EF_{min}$ | -0.76±0.005 | -0.565±0.056 | -0.64±0.012 |
| DA | 2.1±0.084 | 1.44±0.005 | 1.66±0.054 |
| **Femoral midshaft** | | | |
| *Cortical Bone* | | | |
| Mean Ct.Th (mm) | 0.14±0.001 | 0.099±0.002 | 0.13±0.007 |
| BV/TV (%) | 21.72±0.73 | 14.46±0.75 | 16.51±0.44 |
| Ps.Pm (mm) | 4.935±0.08 | 4.36±0.02 | 4.63±0.02 |
| Ec.Pm (mm) | 3.730±0.06 | 3.70±0.07 | 3.66±0.01 |
| J ($mm^4$) | 0.507±0.04 | 0.287±0.026 | 0.355±0.022 |

Data were collected using X-ray Computed Tomography of distal femur and femoral midshaft. Values represent the mean±SEM.

BV/TV= Bone Volume/Total Volume; Tb.N= Trabecular Number; Tb.Th= Trabecular Thickness; Tb.Sp= Trabecular Separation; BS/BV= Specific Bone Surface: Bone Surface Area/Bone Volume; EF= Ellipsoidal Factor; DA= Degree of Anisotropy; Ct.Th= Cortical Thickness; Ps.Pm= Periosteal Perimeter; Ec.Pm= Endocortical Perimeter; J= Polar Moment of Inertia.

**Supplementary Table II**. Summary statistics of each MR regression in the MESA cohort. All statistically significant tests, including intercept tests for vertical pleiotropy, have their associated p-values highlighted in red.

| Method | Estimate | Standard Error | 95% CI lower | 95% CI upper | p-value |
|---|---|---|---|---|---|
| Simple median | 0.033 | 0.267 | -0.49 | 0.556 | 0.9 |
| Weighted median | 0.093 | 0.244 | -0.38 | 0.57 | 0.703 |
| Penalized weighted median | 0.093 | 0.244 | -0.38 | 0.57 | 0.703 |
| IVW | 0.147 | 0.194 | -0.23 | 0.53 | 0.446 |
| Penalized IVW | 0.147 | 0.194 | -0.23 | 0.53 | 0.446 |
| Robust IVW | 0.144 | 0.126 | -0.1 | 0.39 | 0.251 |
| Penalized robust IVW | 0.144 | 0.126 | -0.1 | 0.39 | 0.251 |
| MR-Egger | 0.755 | 0.513 | -0.23 | 1.78 | 0.131 |
| Intercept | -0.101 | 0.076 | -0.25 | 0.049 | 0.186 |
| Penalized MR-Egger | 0.775 | 0.513 | -0.23 | 1.78 | 0.131 |
| Intercept | -0.101 | 0.076 | -0.25 | 0.049 | 0.186 |
| Robust MR-Egger | 0.775 | 0.074 | 0.63 | 0.92 | 0 |
| Intercept | -0.101 | 0.024 | -0.147 | -0.05 | 0.000019 |
| Penalized robust MR-Egger | 0.775 | 0.074 | 0.63 | 0.92 | 0 |
| Intercept | -0.101 | 0.024 | -0.147 | -0.05 | 0.000019 |

**Supplementary Table III**. Summary statistics of each MR regression in the Offspring Cohort of the Framingham Heart Study (FHS). All statistically significant tests, including intercept tests for vertical pleiotropy, have their associated p-values highlighted in red.

| Method | Estimate | Standard Error | 95% CI lower | 95% CI upper | p-value |
|---|---|---|---|---|---|
| Simple median | 0.164 | 0.422 | -0.662 | 0.991 | 0.697 |
| Weighted median | 0.282 | 0.361 | -0.426 | 0.991 | 0.435 |
| Penalized weighted median | 0.282 | 0.361 | -0.426 | 0.991 | 0.435 |
| IVW | 0.235 | 0.312 | -0.376 | 0.847 | 0.451 |
| Penalized IVW | 0.235 | 0.312 | -0.376 | 0.847 | 0.451 |
| Robust IVW | 0.238 | 0.194 | -0.141 | 0.617 | 0.219 |
| Penalized robust IVW | 0.238 | 0.194 | -0.141 | 0.617 | 0.219 |
| MR-Egger | 0.688 | 0.723 | -0.726 | 2.1 | 0.34 |
| Intercept | -0.085 | 0.123 | -0.323 | 0.154 | 0.487 |
| Penalized MR-Egger | 0.688 | 0.723 | -0.726 | 2.1 | 0.34 |
| Intercept | -0.085 | 0.123 | -0.323 | 0.154 | 0.487 |
| Robust MR-Egger | 0.688 | 0.142 | 0.411 | 0.965 | $1.17\times10^{-6}$ |
| Intercept | -0.085 | 0.045 | -0.173 | 0.0036 | 0.06 |
| Penalized robust MR-Egger | 0.688 | 0.142 | 0.411 | 0.965 | $1.17\times10^{-6}$ |
| Intercept | -0.085 | 0.045 | -0.173 | 0.0036 | 0.06 |

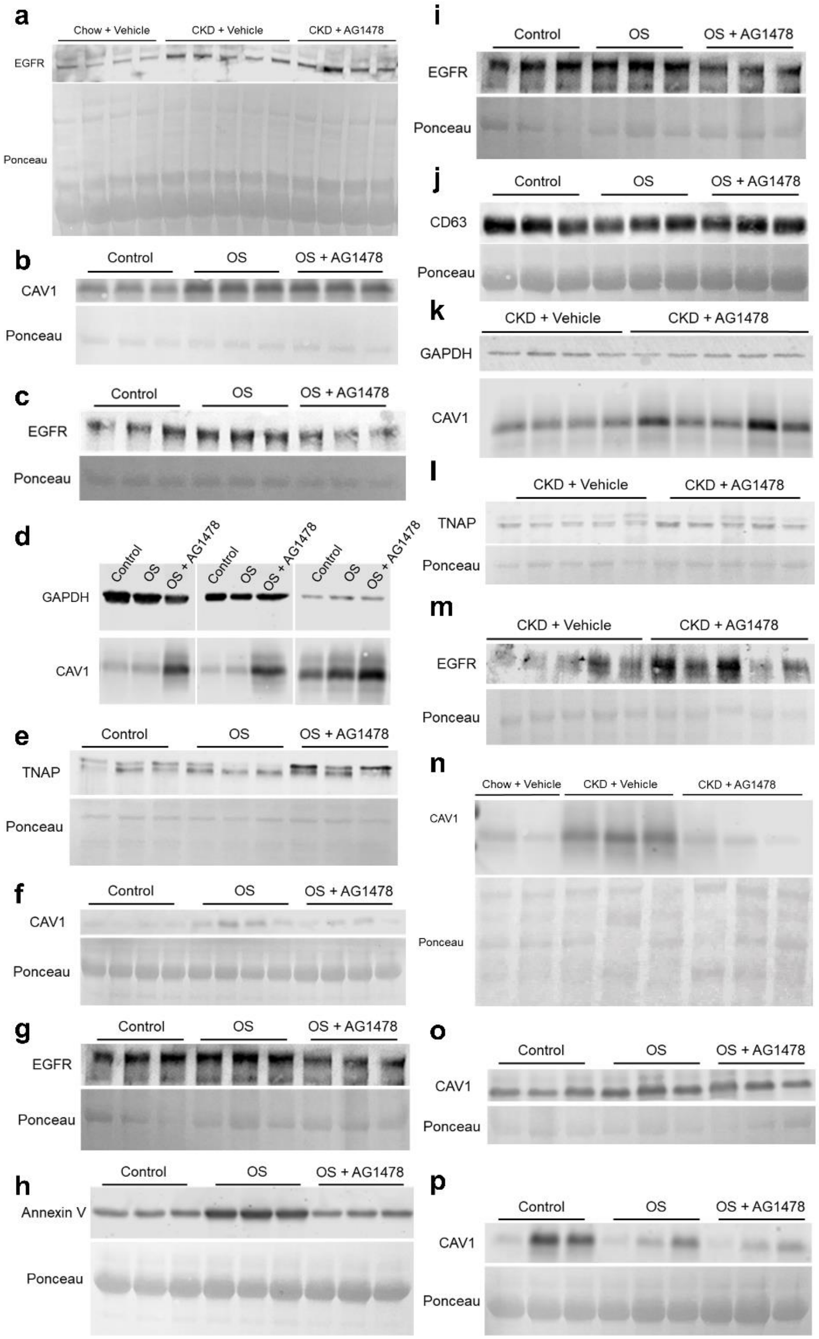


**Supplementary Figure I**. Blot and Ponceau staining for: (A) serum EGFR, (B) intracellular CAV1 in VSMCs, (C) Intracellular EGFR in VSMCs, (D) VSMCs cytosolic CAV1 and GAPDH, (E) VSMCs cytosolic TNAP, (F) VSMCs EV CAV1, (G) VSMCs EV EGFR, (H) VSMCs EV Annexin V, (I) VSMCs EV EGFR, (J) VSMCs EV CD63, (K) Aorta cytosolic CAV1 and GAPDH, (L) Aorta cytosolic TNAP, (M) Aorta cytosolic EGFR, (N) Aorta EV CAV1, (O) Intracellular CAV1 in HOBs, and (P) HOBs Matrix Vesicles CAV1.

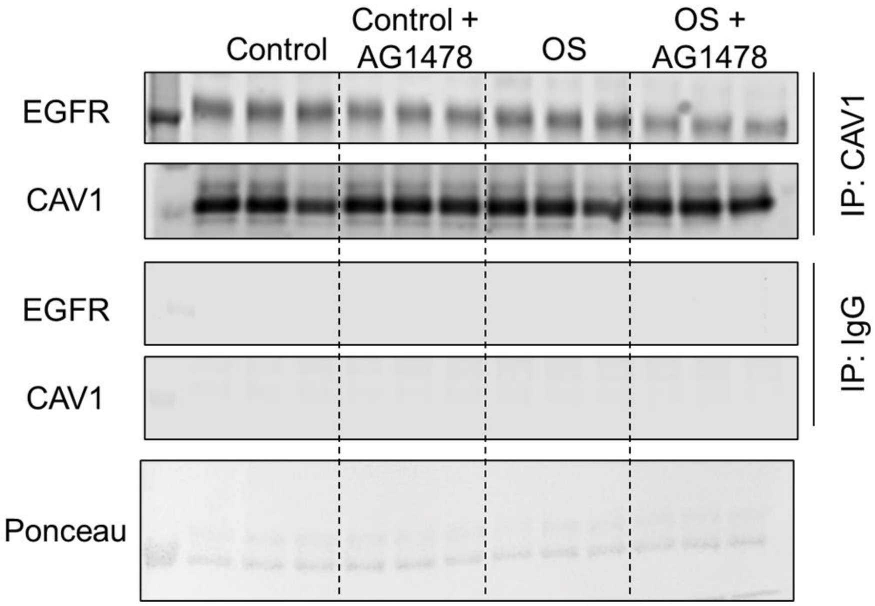


**Supplementary Figure II**. Blot and Ponceau staining for CAV1 and IgG control immunoprecipitation.

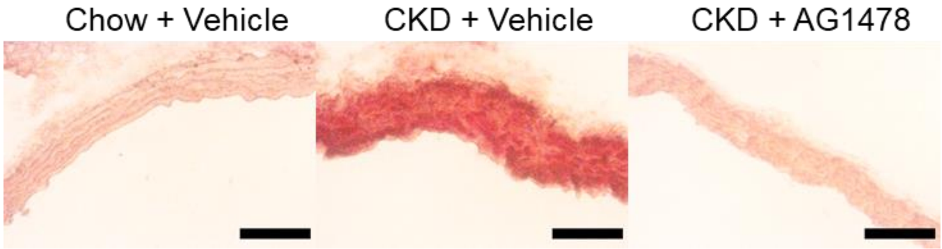


**Supplementary Figure III**. Alizarin Red S staining on the resected aortic tissues from the arch area (10X, scale bar 0.5 mm).
